# Chromatin-associated microprocessor assembly is regulated by PRP40, the U1 snRNP auxiliary protein

**DOI:** 10.1101/2022.01.31.478465

**Authors:** Agata Stepien, Jakub Dolata, Tomasz Gulanicz, Dawid Bielewicz, Mateusz Bajczyk, Dariusz J. Smolinski, Zofia Szweykowska-Kulinska, Artur Jarmolowski

## Abstract

Cotranscriptional processing of RNA polymerase II-generated primary transcripts is a well-documented phenomenon. We recently showed that in plants, miRNA biogenesis is also a cotranscriptional event. Here, we report that Arabidopsis PRP40, the U1 snRNP auxiliary protein, positively regulates the recruitment of SE, the core component of the plant microprocessor, to miRNA genes. The association of DCL1, the microprocessor endoribonuclease, with chromatin was altered in *prp40ab* mutant plants. Impaired cotranscriptional microprocessor assembly was accompanied by RNA polymerase II accumulation at miRNA genes and retention of miRNA precursors at their transcription sites in the *prp40ab* mutant plants. We show that cotranscriptional microprocessor assembly, regulated by AtPRP40, positively affects RNAPII transcription of miRNA genes and is important to reach the correct levels of produced miRNAs.

## Introduction

microRNAs (miRNAs) are single-stranded, usually 21 nt long, RNAs that regulate basic developmental processes as well as plant responses to constant fluctuations in environmental conditions ^1–4^. Therefore, miRNA production has to be tightly controlled at multiple levels ^5,6^. Plant miRNA genes (*MIR*s) are transcribed by RNA polymerase II (RNAPII) to generate primary precursors (pri-miRNAs) that are cleaved to pre-miRNAs (hairpin structures) and further to miRNA/miRNA* duplexes. In plants, both steps of miRNA biogenesis are carried out in the nucleus by the RNase III-type ribonuclease DCL1 (Dicer-like 1) ^7^. Depending on the pri-miRNA structure, DCL1 generates the first cut near the base of the hairpin structure (base-to-loop processing, BTL) or starts from the hairpin loop (loop-to-base processing, LTB) ^8,9^. DCL1 is assisted by SE (SERRATE) and HYL1 (HYPONASTIC LEAVES 1) for accurate and efficient activity ^10–12^. These three proteins, DCL1, SE and HYL1, form the core of the plant microprocessor.

Similar to that of animals, plant pri-miRNA cleavage was recently reported to occur cotranscriptionally ^13,14^. Two components of the microprocessor, DCL1 and SE, are known to be associated with chromatin. The association of DCL1 with *MIR*s is mediated by the Mediator and Elongator complexes ^15,16^, while SE was found to bind and regulate the transcription of intronless genes^17^. However, the cotranscriptional mechanism of the microprocessor assembly is still unclear and requires investigation.

We previously showed that the SE and U1 snRNP proteins are responsible for the communication between the microprocessor and the splicing machinery ^18^. Among the U1 snRNP partners of SE, PRP40 is particularly interesting. Since U1 snRNP is recruited cotranscriptionally to intron-containing genes in yeast shortly downstream from the 5’ splice site, U1 snRNP elements have been considered potential players in the well-established crosstalk between the splicing and transcription machinery ^19–25^. The reported direct interaction of yeast PRP40p with the RNAPII C-terminal domain (CTD) ^26^ indicates that this protein is a good candidate for such communication. Plant and animal PRP40 homologs can also interact with CTD ^27–29^. However, the published data suggest that the main role of PRP40 is stabilization of U1 snRNP by the interactions of PRP40 with other U1 components and modulation of alternative splicing ^26,30–34^, and its possible function in the cotranscriptional recruitment of different macromolecular complexes to RNAPII is still an open question.

We previously suggested that PRP40 may be involved in cotranscriptional microprocessor assembly due to the direct interaction between AtPRP40 and SE ^18^. Here, we found that the levels of almost half of the polyadenylated pri-miRNAs were affected in *prp40ab* (the majority of which were downregulated). However, in parallel, we observed increased levels of chromatin-associated, non-poly(A) tailed transcripts. Interestingly, mature miRNAs, on average, showed only slightly upregulated expression in *prp40ab*. We demonstrate that the higher accumulation of RNAPII along pre-miRNA genes correlates with the altered recruitment of microprocessor components, e.g., SE and DCL1, to *MIR*s. Our results show that in plants, PRP40 is involved in the coordination of the microprocessor assembly on pri-miRNAs while they are synthesized by RNAPII.

## Results

### AtPRP40 is important for *Arabidopsis thaliana* development

The *A. thaliana* genome encodes three PRP40 genes ^35^ that show differences in expression levels in various tissues and developmental stages (Extended Data Fig. 1). AtPRP40a showed the highest expression, whereas AtPRP40c was barely expressed in all tested samples. In our previous studies, we identified AtPRP40a and AtPRP40b as proteins involved in the interplay between miRNA production and the splicing of miRNA precursors ^18^. Interestingly, when we assessed the colocalization of AtPRP40b and U1 snRNA, a core component of U1 snRNP, we found low colocalization coefficients (Extended Data Fig. 2). This finding suggests an additional activity of AtPRP40 beyond the U1 snRNP complex.

Single mutants of each AtPRP40 did not differ from WT plants (Extended Data Fig. 3a). However, the double *prp40ab* mutant showed delayed growth (Extended Data Fig. 3b, c). Interestingly, the AtPRP40a mRNA level in the single *prp40b* mutant was elevated compared to that in the WT plants, and similarly, *AtPRP40b* mRNA expression was higher in the *prp40a* mutant than in the WT plants (Extended Data Fig. 4a). This result was also confirmed at the protein level using antibodies recognizing AtPRP40b (Extended Data Fig. 4b). Notably, the level of AtPRP40c mRNA was not affected in *prp40a, prp40b* or *prp40ab* (Extended Data Fig. 4a). Since we were not able to obtain homozygotes of the triple *prp40abc* mutant (most likely due to a relatively short distance between the *AtPRP40B* and *AtPRP40C* genes, approximately 54 kb), for further analyses, we used the double *prp40ab* mutant. Interestingly, we found that the crosstalk between SE and AtPRP40a and AtPRP40b is also crucial for plant development, as the mutation of *SE* and inactivation of *AtPRP40A* and *AtPRP40B* led to embryo lethality (Extended Data Fig. 5). Notably, we did not observe this phenomenon after crossing *prp40ab* with *hyl1-2*, a mutant of another core microprocessor component, HYL1. This finding prompted us to analyze the crosstalk between the SE and AtPRP40 proteins and its role in miRNA biogenesis in plants.

### AtPRP40 mediates the SE association with RNAPII and *MIR*s

Since SE was shown to associate with RNAPII ^17^, we investigated whether this contact is regulated by AtPRP40. We performed colocalization analyses and utilized the proximity-ligation assay (PLA). These experiments showed the close proximity and colocalization of SE and AtPRP40b in the plant cells (Fig. 1a, Extended Data Fig. 6), and the close proximity and colocalization of SE with RNAPII phosphorylated at both Ser5 (P-Ser5-RNAPII) and Ser2 (P-Ser2-RNAPII) (Fig. 1b, Extended Data Fig. 7). The colocalization coefficients calculated for SE and both phosphorylated forms of RNAPII were significantly lower in the *prp40ab* mutant plants (Extended Data Fig. 7b), and the PLA signal numbers also decreased in *prp40ab* (Fig. 1b), indicating that AtPRP40 mediates the association of SE with RNAPII.

**Fig 1.**
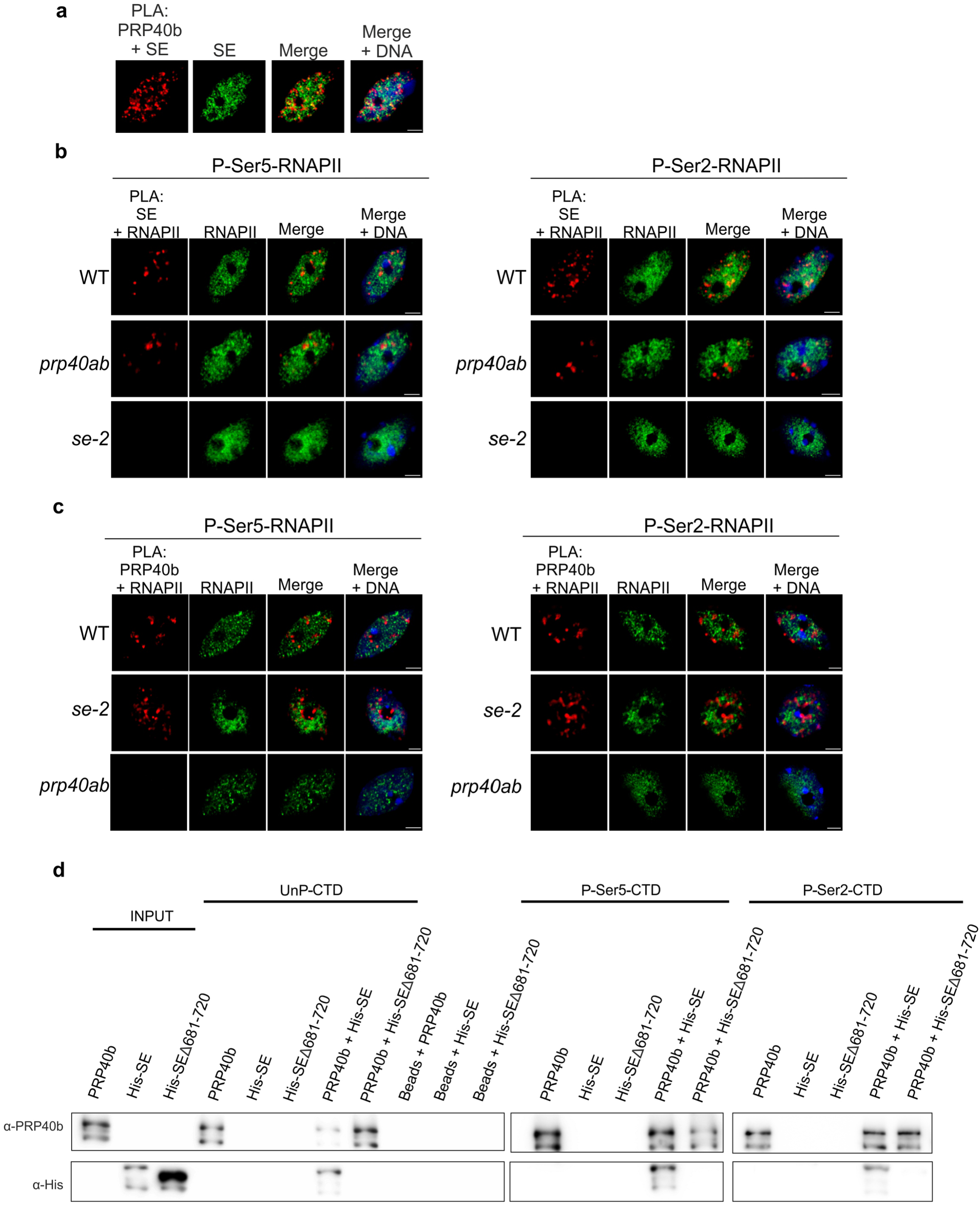
AtPRP40 regulates the association of SE with RNAPII. **a**, Close proximity of AtPRP40b and SE in the cell nucleus (first image, red signals) analyzed by PLA in view of SE nuclear localization (second image, green signals). DNA was stained with Hoechst (blue). Scale bar = 5 µm. **b**, Close proximity of SE and RNAPII phosphorylated at CTD Ser5 or Ser2 in the WT, *prp40ab* and *se-2* plants (first columns, red signals) in view of P-Ser5-RNAPII or P-Ser2-RNAPII nuclear localization (second columns, green signals) detected by PLA. DNA was stained with Hoechst (blue). Scale bar = 5 µm. **c**, Close proximity of AtPRP40b and RNAPII phosphorylated at CTD Ser5 or Ser2 in the WT, *se-2* and *prp40ab* plants (first columns, red signals) in view of P-Ser5-RNAPII or P-Ser2-RNAPII nuclear localization (second columns, green signals) detected by PLA. DNA was stained with Hoechst (blue). Scale bar = 5 µm. **d**, *In vitro* pull-down assays using recombinant AtPRP40b and SE proteins (both full length and the shortened C-terminus variant b.681-720 were used) and CTD peptides in unphosphorylated (UnP) or phosphorylated Ser5 (P-Ser5-CTD) or Ser2 (P-Ser2-CTD) forms. Input represents 1/10 of the protein sample.

In contrast, the colocalization and close proximity of AtPRP40b with RNAPII do not depend on SE, since we did not observe any change in the colocalization coefficient values or the observed PLA signals in *se-2* (Extended Data Fig. 8, Fig. 1c).

Moreover, using an *in vitro* pull-down assay we showed that AtPRP40b is required for the association of SE with the C-terminal domain (CTD) of both the unphosphorylated and phosphorylated forms of RNAPII (Fig. 1d). In this experiment, we also generated the SE protein lacking forty C-terminal amino acids (SEΔ681-720) that corresponds to the SE variant expressed in the *se-2* mutant. We previously showed that such a truncated SE protein cannot bind AtPRP40b ^18^; thus, we used this SE variant as a negative control. Indeed, we observed SE-AtPRP40b-CTD complex formation only in the presence of the full-length SE but not when the SEΔ681-720 shortened variant of SE was used (Fig. 1d).

Furthermore, we performed a ChIP experiment to determine whether AtPRP40 is involved in SE recruitment to miRNA genes, and we found lower SE accumulation on all *MIR*s tested in *prp40ab* than in the wild-type plants (Fig. 2, Extended Data Fig. 9).

**Fig. 2.**
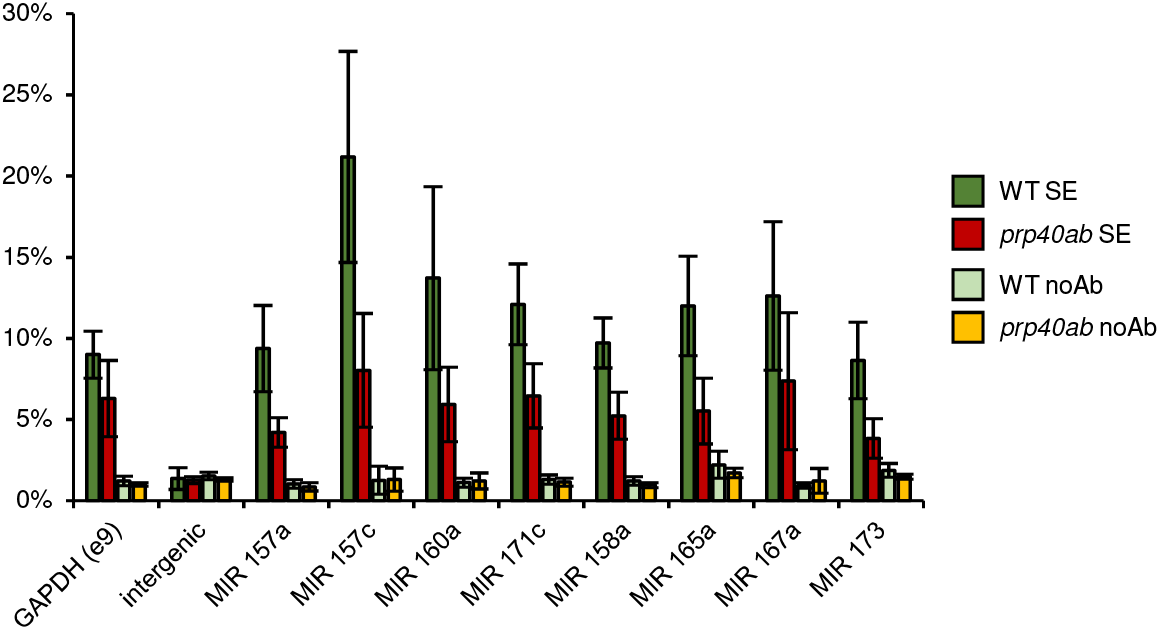
AtPRP40 is required for the proper accumulation of SE on miRNA genes. Quantitative ChIP-PCR analyses of SE accumulation on randomly selected miRNA genes in the *prp40ab* mutant (n=3).

Our data indicate that AtPRP40 regulates the recruitment of SE to RNAPII and *MIR*s and may be involved in the regulation of miRNA biogenesis.

### AtPRP40 is involved in the transcription of pri-miRNAs

To test the role of AtPRP40 in miRNA production, we applied our high-throughput RT–qPCR platform, mirEX 2.0, which allowed us to analyze the expression levels of 297 *A. thaliana* pri-miRNAs ^36,37^. The levels of 46% polyadenylated pri-miRNAs were significantly changed in *prp40ab*, (121 out of 261 pri-miRNAs which we were able to detect) (Fig. 3a). The vast majority (71%) of affected precursors showed downregulated expression. The affected pri-miRNAs belong to both the low- and high-expression pri-miRNAs; thus, AtPRP40 protein activity does not depend on the pri-miRNA expression level (Extended Data Fig. 10a). Moreover, we did not find any relationship between the expression pattern in *prp40ab* and the presence of introns in *MIR* genes (Extended Data Fig. 10b). To determine how this decreased level of polyadenylated pri-miRNAs in *prp40ab* mutants affects mature miRNAs, we performed small RNA sequencing and compared the miRNA levels of the WT and *prp40ab* plants. Surprisingly, the levels of most of the mature miRNAs were not changed or were even slightly upregulated in the *prp40ab* mutants (Fig. 3b). The affected miRNAs were mostly highly expressed miRNAs (Extended Data Fig. 10c). Interestingly, the miRNAs showing no change or upregulated expression correspond mainly to those poly(A) pri-miRNAs that showed lower levels in *prp40ab* (Fig. 3c, d). Furthermore, we investigated whether precision of miRNA production is impaired in *prp40ab*, and we did not observe any bias in the *prp40ab* mutants compared to the WT plants (Extended Data Fig. 10d). To elucidate this unexpected phenomenon, we randomly selected a group of *MIR*s (including polyadenylated *MIR* transcripts with upregulated, downregulated and unaffected levels) and tested their expression levels using cDNA templates synthesized with random hexamer primers instead of oligo d(T), which was used to monitor the levels of poly(A)-tailed pri-miRNAs. Interestingly, none of these molecules were significantly changed (Extended Data Fig. 10e). We wanted to exclude the possibility that the effect on poly(A)-tailed pri-miRNA levels was due to the change in the amount of DCL1, a major component of the microprocessor. To this end we performed the western blot analysis in WT and the *prp40ab* plants.The levels of DCL1 and SE, a second key subunit of the microprocessor, were not altered in *prp40ab* (Extended Data Fig. 11). However, we observed an increased level of the double-stranded RNA-binding protein HYL1 in *prp40ab*.

**Fig. 3.**
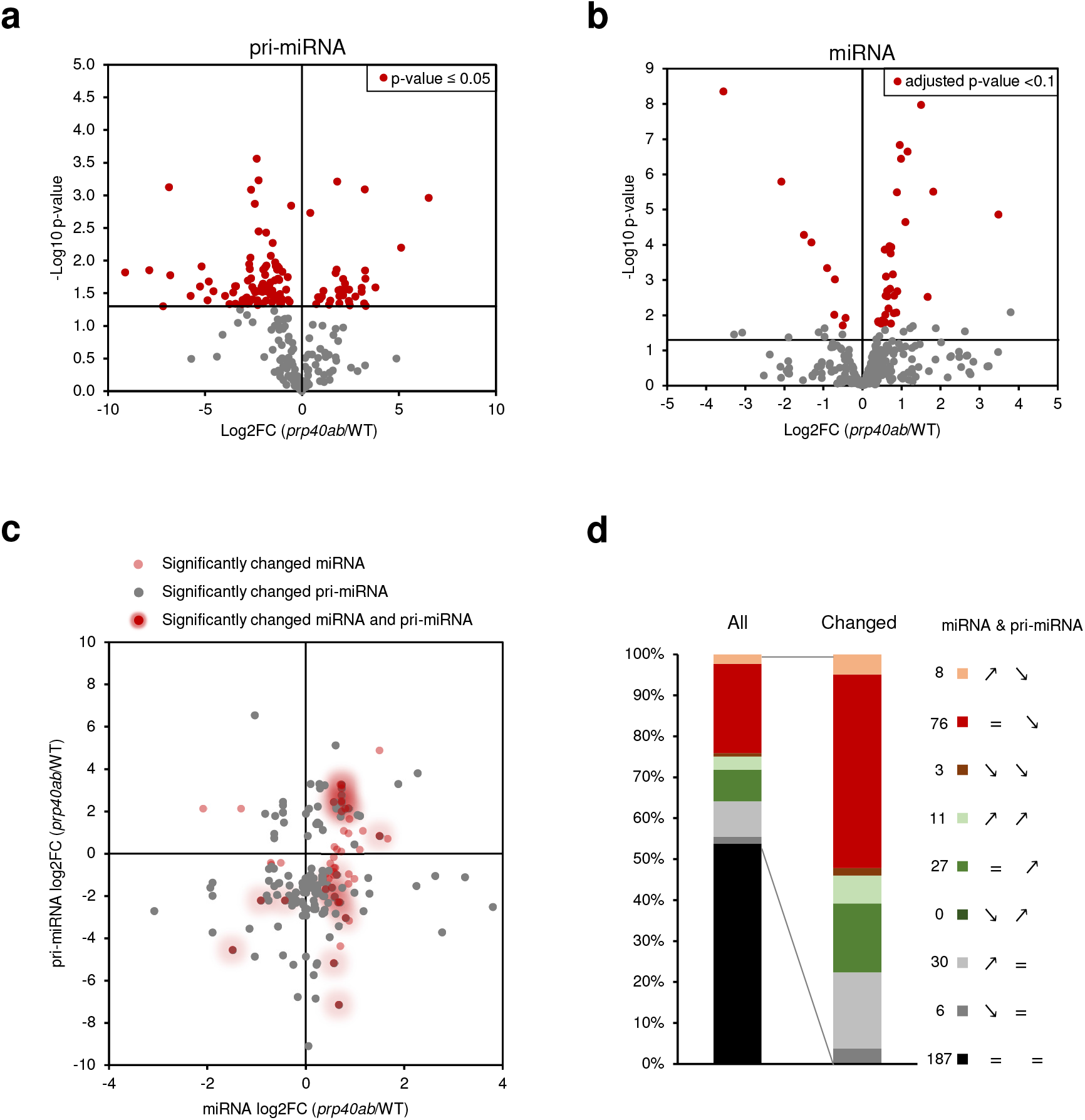
AtPRP40 affects the early steps of miRNA biogenesis. **a**, Volcano plot showing changes in the levels of polyadenylated *MIR* transcripts in *prp40ab* (RT-qPCR, n=3). **b**, Volcano plot showing changes in the levels of miRNAs in *prp40ab* (small RNA sequencing, n=3). **c**, Scatter plot showing the levels of polyadenylated *MIR* transcripts relative to corresponding miRNAs. **d**, Expression patterns in *prp40ab* for polyadenylated *MIR* transcripts paired with the corresponding miRNAs. Numbers on the legend corresponds to miRNA & pri-miRNA pairs from each group. Signs on the legend indicate for expression pattern in *prp40ab* for miRNA and pri-miRNA respectively (↗increased level, ↘decreased level, = not changed).

Thus, these results suggest that the lack of Arabidopsis PRP40a and b affects the accumulation of polyadenylated pri-miRNAs, but the final effect on the production of mature miRNA in *prp40ab* mutants is rather minor.

### AtPRP40 affects RNAPII and DCL1 occupancy on *MIR* genes

The discordance between the levels of poly(A)-tailed pri-miRNAs and the total amount of *MIR* transcripts might be due to improper transcription termination and/or changes in miRNA precursor processing. Whole-genome RNAPII profiling in the *prp40ab* mutant revealed a higher level of RNAPII on *MIR*s with a significant increase in pre-miRNA coding regions and further in 3’ ends (Fig. 4a, b, Extended Data Fig. 12a, c). Additionally, the accumulation of RNAPII along miRNA genes in *prp40ab* in the vast majority of cases correlated with the reduced level of polyadenylated pri-miRNAs (Fig. 4c), and the pri-miRNA coding regions with increased RNAPII occupancy had significantly lower levels of the corresponding poly(A) pri-miRNAs (Fig. 4d).

**Fig. 4.**
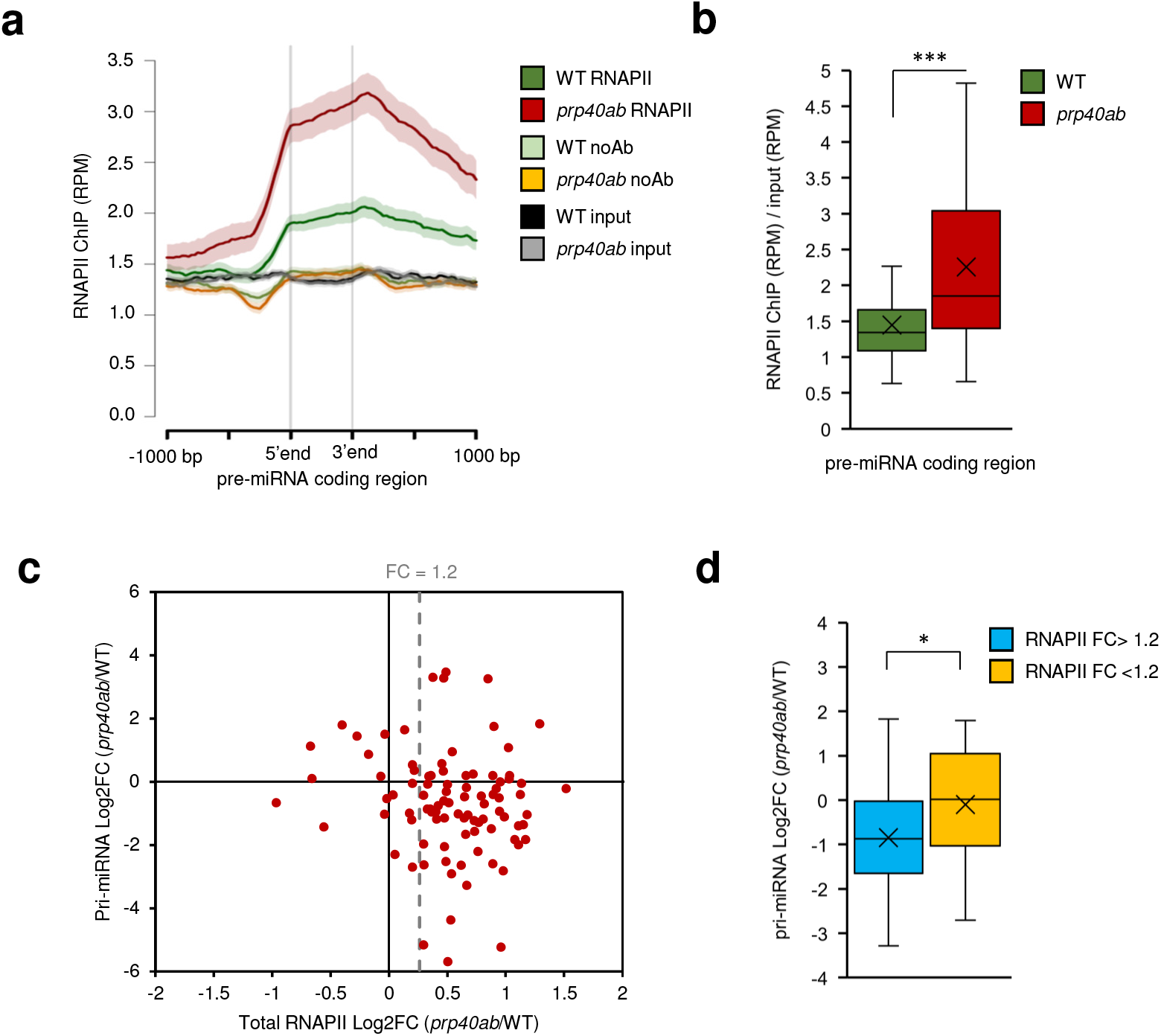
RNAPII distribution on miRNA genes is affected in the *prp40ab* mutant. **a**, Metagene analysis of RNAPII distribution on pre-miRNA coding regions based on ChIPseq data **b**, Box plot showing the RNAPII occupancy on pre-miRNA coding regions based on ChIPseq data. **c**, Scatter plot showing changes in RNAPII occupancy on pre-miRNA coding regions relative to changes in the levels of polyadenylated *MIR* transcripts in *prp40ab*. **d**, Box plot showing changes in the levels of polyadenylated *MIR* transcripts in *prp40ab* depending on a change in the RNAPII level in pre-miRNA coding regions. Mann–Whitney U test p value: * < 0.05; *** < 0.001. The box is drawn between the first and third quartiles, with an additional line drawn along the second quartile to mark the median. “X” indicates for mean. Whiskers indicate the minimums and maximums outside the first and third quartiles. The shaded area around each curve on metaplots indicates standard errors.

Previously, DCL1 was detected on at least some *MIR*s ^16^; thus, we investigated how the lack of the AtPRP40a and b proteins affects DCL1 accumulation on chromatin. We observed alteration of the DCL1 level in pre-miRNA coding regions (Fig. 5a, Extended Data Fig. 12b, c). On average, DCL1 occupancy was increased; however, there was a group of *MIR*s with unchanged or even decreased DCL1 levels. We recently showed that cotranscriptional processing of pri-miRNAs differs depending on the way precursors are cleaved at the first step ^14^. For the loop-to-base (LTB) type of cleavage, both processing steps occur cotranscriptionally, whereas for the base-to-loop (BTL) type of processing, the first step is cotranscriptional, but the second occurs post-transcriptionally in the nucleoplasm. We found that LTB-type *MIR*s, but not BTL-type *MIR*s, showed increased DCL1 occupancy in *prp40ab* (Fig. 5b, c).

**Fig. 5.**
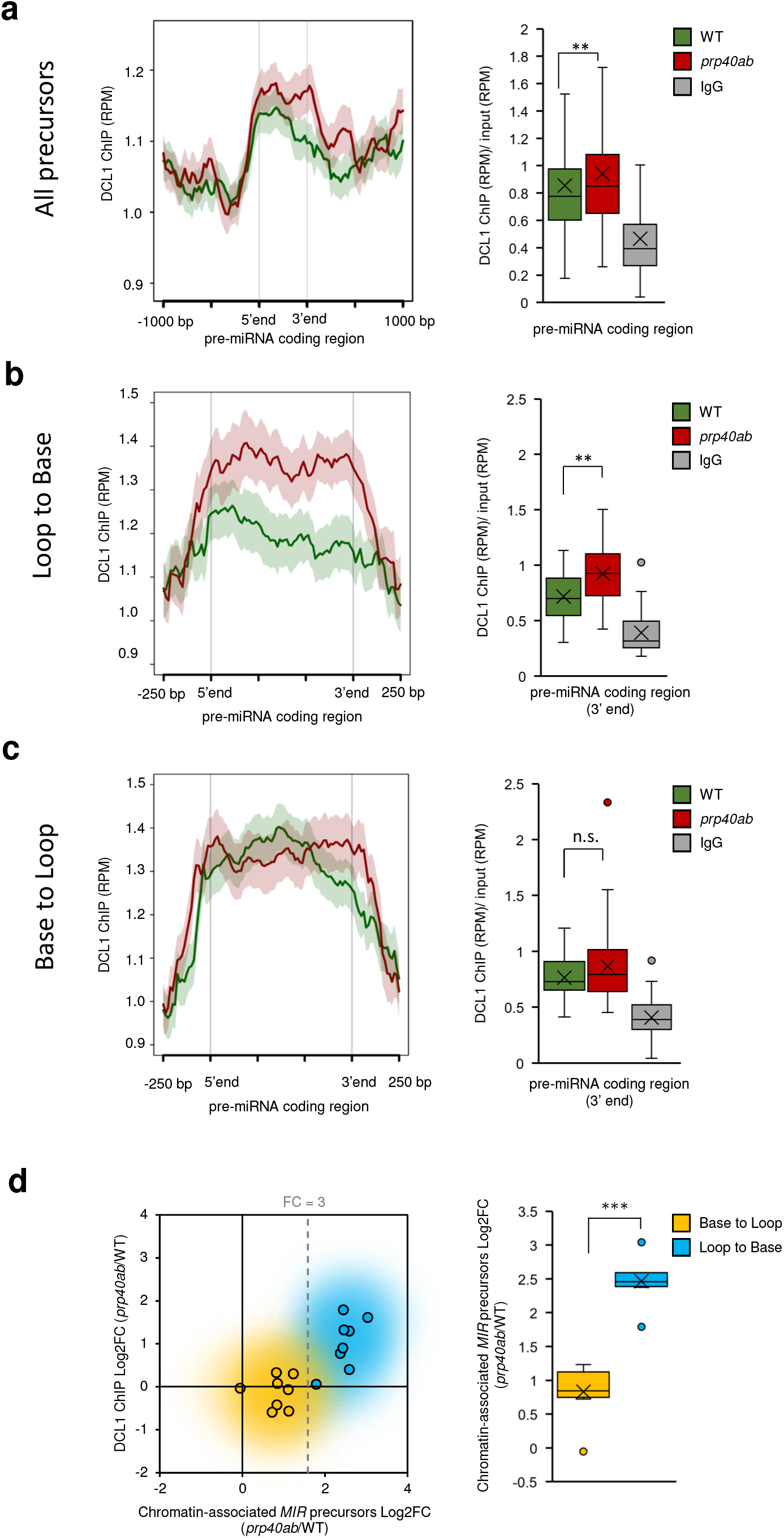
DCL1 distribution on miRNA genes is affected in the *prp40ab* mutant. **a**, DCL1 distribution on pre-miRNA coding regions based on ChIPseq data. **b**, DCL1 distribution on loop-to-base-type miRNA genes. **c**, DCL1 distribution on base-to-loop-type miRNA genes. **d**, The level of chromatin-associated *MIR* transcripts relative to the level of DCL1 on pre-miRNA coding regions depending on the miRNA gene type. Mann–Whitney U test p value: ** < 0.01; *** < 0.001. The box is drawn between the first and third quartiles, with an additional line drawn along the second quartile to mark the median. “X” indicates for mean. Whiskers indicate the minimums and maximums outside the first and third quartiles. The shaded area around each curve on metaplots indicates standard errors.

Both RNAPII and DCL1 ChIP-seq datasets strongly indicate that AtPRP40 proteins are involved in the cotranscriptional regulation of *MIR* expression.

The increased RNAPII and/or DCL1 occupancy on pre-miRNA coding regions suggests that in *prp40ab*, transcription and processing complexes are stuck on *MIR*s. To determine whether miRNA precursors also accumulated on *MIRs* in the *prp40ab* mutants compared to the WT plants, we separated the nucleoplasmic and chromatin fractions of nuclei (Extended Data Fig. 13a) and tested the pri-miRNA levels with RT-qPCR performed separately for each fraction. In agreement with our previous data ^14^, we found that in the WT, the pre-miRNAs derived from BTL-type precursors accumulated predominantly in the nucleoplasm (Extended Data Fig. 13b). Furthermore, we calculated the *prp40ab*/WT fold change for chromatin and nucleoplasmic fractions and found that LTB-type precursors are highly associated with chromatin in *prp40ab* (fold change over 3) (Fig. 5d, Extended Data Fig. 13c), while BTL-type precursors show only minor (fold change below 3) or no change in the chromatin fraction (Fig. 5d, Extended Data Fig. 13d). In the nucleoplasmic fraction, both types of precursors were mostly decreased. Interestingly, we did not find an increased chromatin association of full-length poly(A)-tailed precursors (Extended Data Fig. 13e).

The obtained results show that AtPRP40 regulates cotranscriptional miRNA biogenesis by affecting RNAPII and microprocessor activity on *MIR*s. This finding shows that AtPRP40 is involved in correct cotranscriptional microprocessor assembly.

## Discussion

Coupling of pre-mRNA processing with transcription is a well-established phenomenon ^23–25,38–40^. Cotranscriptional processing of miRNA precursors in humans has also been reported ^13,41^. However, some results obtained demonstrate that mammalian pri-miRNA processing kinetics range from fast over intermediate to slow and that pri-miRNAs might be processed both co- and post-transcriptionally ^42^. Moreover, it has been claimed that chromatin retention does not determine the processing type, since the authors observed chromatin release of pri-miRNAs at comparable times after transcription for pri-miRNAs showing different processing kinetics. However, the factors regulating the kinetics of pri-miRNA processing and the co- and post-transcriptional mechanism of pri-miRNA processing have not been identified.

In human cells, the FUS (fused in sarcoma/translocated in liposarcoma) protein has been suggested to be a factor involved in the biogenesis of a large class of miRNAs, among which neuronal miRNAs are known to have a crucial role in neuronal function ^43^. It has been shown that FUS is recruited to chromatin by binding to newly synthetized pri-miRNAs, where it facilitates Drosha (one of the two RNase III enzymes involved in miRNA biogenesis in animals) loading, supporting cotranscriptional processing of miRNA precursors ^43^. It has also been suggested that the histone H1-like protein HP1BP3, but not canonical H1 variants, associates with the human microprocessor and promotes global miRNA biogenesis. HP1BP3 binds both DNA and pri-miRNA and enhances cotranscriptional miRNA processing *via* chromatin retention of nascent pri-miRNAs. This study clearly suggests the existence of a class of chromatin retention factors stimulating cotranscriptional miRNA processing in animal cells ^44^. Thus, the results indicate that both specific chromatin marks and additional protein factors connected with miRNA gene transcription may control the cotranscriptional assembly of the miRNA biogenesis machinery.

In contrast to that in animals, plant miRNA biogenesis occurs exclusively in the cell nucleus. Moreover, special nuclear structures called dicing bodies (D-bodies) are considered places of plant pri-miRNA processing ^45^. The fact that in each plant cell nucleus, only a few D-bodies are observed may suggest a post-transcriptional mechanism of plant pri-miRNA processing. Recently, we showed that pri-miRNA processing in plants is a cotranscriptional process; however, some steps occur post-transcriptionally in the nucleoplasm after release of the processing intermediates (pre-miRNAs) from the transcription sites. Moreover, we discovered that the structure of miRNA primary precursors dictates processing localization ^14^. Plant pri-miRNAs were shown to be processed in two different manners: base-to-loop (BTL processing) and loop-to-base (LTB processing) ^8,9^. We demonstrated that for pri-miRNAs processed in an LTB manner, both processing steps occur cotranscriptionally, and in the case of BTL-type pri-miRNA processing, the first step is cotranscriptional, but the second occurs post-transcriptionally in the nucleoplasm. The data presented in this work confirm our previous conclusions on cotranscriptional miRNA biogenesis in plants ^14^. We show here that BTL-type miRNA precursors accumulate predominantly in the nucleoplasm (Extended Data Fig. 13b), in contrast to LTB-type transcripts that localize mostly on *MIR*s. In this paper, we also identified a protein factor that is involved in the regulation of cotranscriptional miRNA biogenesis in plants. This protein is AtPRP40, the Arabidopsis U1 snRNP auxiliary protein.

The direct interaction between AtPRP40 that binds to the CTD of RNAPII and the SE protein has been described by us previously ^18^. This observation prompted us to test whether SE forms a complex with RNAPII and whether the SE/RNAPII interaction requires the presence of AtPRP40. Indeed, we prove here that AtPRP40 mediates the association of SE with RNAPII (Fig. 1b, d, Extended Data Fig. 7) on miRNA genes (Fig. 2). Therefore, we further explored the role of AtPRP40 in miRNA biogenesis and cotranscriptional microprocessor assembly. We observed lower levels of polyadenylated miRNA precursors (Fig. 3a) but increased accumulation of non-polyadenylated precursors on *MIR*s in the *prp40ab* mutant plants (Extended Data Fig. 13). We also found that RNAPII accumulates in the *prp40ab* mutants on pre-miRNA coding regions, which together indicate retention of RNAPII on *MIR*s when the AtPRP40a and b proteins are absent (Fig. 4, Extended Data Fig. 12a, c). The recruitment of DCL1 and SE, two key components of the plant microprocessor complex, to *MIR*s is also affected in *prp40ab*, suggesting a role of AtPRP40 in cotranscriptional microprocessor assembly (Figs. 2, 5, Extended Data Figs. 9, 12b, c). Moreover, our results indicate the existence of an interplay between the microprocessor and RNAPII. The influence of pre-mRNA processing on RNAPII activity was observed previously in the case of splicing. It has been shown that inactivation of the promoter proximal 5′ splice sites reduces the level of nascent transcription ^46^. Recently, it has also been reported that U2 snRNP has a positive effect on RNAPII transcription elongation ^47^. The existence of a transcriptional elongation checkpoint that is associated with cotranscriptional presplicosome formation has been previously reported by the Beggs group ^48^. In Arabidopsis, we previously showed that NTR1, a spliceosome disassembly factor, is responsible for slowing RNAPII and the formation of splicing checkpoints at alternative splice sites ^49^. Thus, similar to the interplay between the splicing and transcription machinery, we postulate here that correct microprocessor assembly, regulated by AtPRP40, has a positive effect on RNAPII transcription. In WT plants, AtPRP40 mediates the recruitment of SE to nascent *MIR* transcripts and facilitates the proper assembly of the whole microprocessor, processing of the primary miRNA precursors (Fig. 6) and the smooth movement of RNAPII along pre-miRNA coding regions. In the case of impaired microprocessor component recruitment, as we show in the *prp40ab* mutants, RNAPII slows down to allow the miRNA biogenesis complex to be formed.

**Fig. 6.**
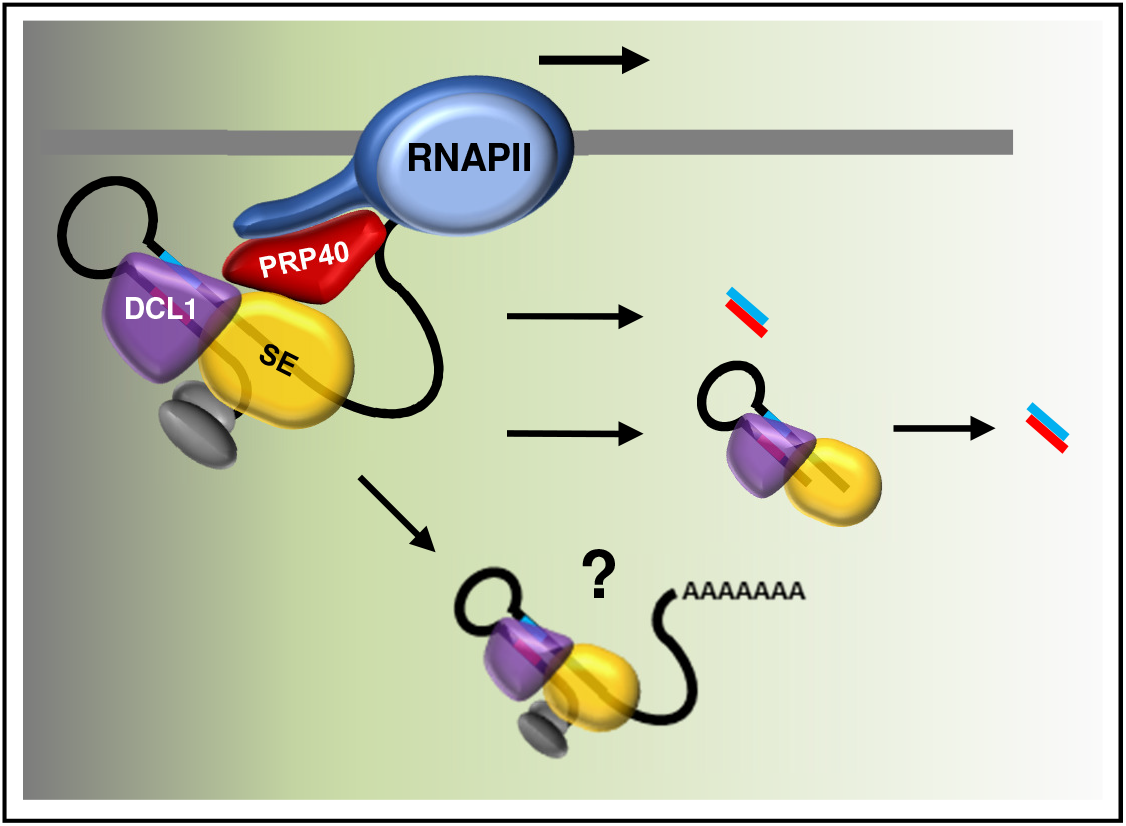
Cotranscriptional microprocessor assembly, regulated by AtPRP40, impacts RNAPII activity and is required for correct miRNA production. In WT plants, when AtPRP40 is present, SE and DCL1 are recruited to miRNA genes, primary precursors are efficiently processed to pre-miRNAs (hairpin structures) and further to miRNAs, and RNAPII fluently moves through the pre-miRNA coding region. In the case of loop-to-base-type miRNA genes, both processing steps occur cotranscriptionally, and the miRNA/miRNA* duplex is released to the nucleoplasm. For base-to-loop-type miRNA genes, only the first step is cotranscriptional, and the second cleavage step takes place in the nucleoplasm. Whether the polyadenylated pri-miRNAs that are released from chromatin can be post-transcriptionally processed needs to be verified.

Interestingly, while RNAPII accumulates on most *MIR*s in *prp40ab*, we observed increased occupancy of DCL1 only on approximately half of the *MIR*s in this mutant (Extended Data Fig. 12c). On the rest of the *MIR*s, the accumulation of DCL1 in *prp40ab* was not changed or decreased. DCL1 accumulation on *MIR*s in the *prp40ab* mutant is most likely due to the retention of nascent non-polyadenylated transcripts attached to the transcription sites. As already mentioned, both steps of processing of LTB pri-miRNAs are carried out cotranscriptionally, in contrast to BTL pri-miRNAs, where the second step of miRNA maturation takes place post-transcriptionally after releasing pre-miRNAs to the nucleoplasm. This difference in miRNA biogenesis can explain why the accumulation of DCL1 is observed only on LTB *MIR*s: stem–loop structures are cut out from BTL pri-miRNAs and released to the nucleoplasm, taking DCL1 away from the transcription sites. However, the higher association of DCL1 with *MIR*s and the retention of primary miRNA precursors on chromatin had no major effect on the levels of mature miRNAs (Fig. 3b). This finding indicates that cotranscriptional pri-miRNA processing is very efficient, and in the case of a disturbance in microprocessor assembly, the retention of miRNA precursors at transcription sites is sufficient to obtain the correct levels of mature miRNAs. This finding is in agreement with data showing that animal pri-miRNAs retained at transcription sites due to the deletion of 3’ end processing signals are processed more efficiently than pri-miRNAs that are cleaved, polyadenylated and released ^50^. One of the possible reasons for this result is the higher amount of substrates available for processing. We observed a similar effect of mutation of HEN2, an RNA helicase that is involved in RNA degradation by the nuclear RNA exosome, on miRNA biogenesis ^51^. In *se-2*, pri-miRNAs accumulate, and mature miRNA levels are downregulated because of poor pri-miRNA processing efficiency in the absence of fully active SE. However, in the *se-2 hen2-2* double mutant, we observed the increased accumulation of pri-miRNAs and partially restored levels of mature miRNAs, which led to attenuation of the developmental defects characteristic of *se-2* mutant plants ^51^. This result indicates that slow degradation of pri-miRNAs due to HEN2 loss can compensate for inefficient pri-miRNA processing. In WT plants, cotranscriptional microprocessor assembly on newly synthetized pri-miRNAs stimulated RNAPII to pause the transcription of *MIR*s (Fig. 4a). Impaired cotranscriptional microprocessor assembly leads to longer RNAPII pausing in the pre-miRNA coding region accompanied by the accumulation of miRNA primary precursors at their transcription sites. This phenomenon provides additional time for the processing of accumulated, non-polyadenylated *MIR* transcripts. Therefore, the level of most miRNAs was not changed or was even slightly increased, while the levels of polyadenylated pri-miRNAs decreased in plants lacking AtPRP40a and b. Thus, our results demonstrate that AtPRP40 is the protein that contributes to the cotranscriptional recruitment of DCL1 and SE to pri-miRNAs, which regulate RNAPII activity over *MIR*s. However, we still do not exclude the possibility of a more direct influence of AtPRP40 on transcription carried out by RNAPII. Additional studies are needed to distinguish between these possibilities.

## Methods

### Plant material

*A. thaliana* (Col-0 wild-type, *se-1* ^52^, *se-2* ^53^, *prp40a* (SALK_021070), *prp40b* (SALK_066044), *prp40c* (SALK_148319), *prp40ab* (SALK_021070 x SALK_066044), and *hyl1-2* ^54^ seeds after 3 days of stratification were grown at 22 °C (16/8 h light/dark, 50–60% humidity, 150–200 µmolm-2 s-1 photon flux density) in Jiffy-7 pots (Jiffy) and collected on Day 21 of growth or used in crosses. The GFP-SE transgenic line was produced in the *se-4* ^55^ background by incorporation of an additional copy of SE fused with GFP under the control of the UBQ10 promoter. For ChIP analyses, 14-day-old seedlings grown on ½ MS solid medium were used instead.

### RNA isolation and cDNA preparation

Total RNA for AtPRP40 mRNAs, pri-miRNA and miRNA level analyses was prepared according to ^56^. Briefly, RNA was isolated using a Direct-zol™ RNA Mini Prep Kit (Zymo Research) and treated with Turbo DNase I (Thermo Fisher Scientific).

For transcript expression levels, 3 µg of total DNase-treated RNA was reverse transcribed to cDNA with the use of SuperScript III Reverse Transcriptase (Thermo Fisher Scientific) and oligo-dT(18) or random-hexamer primers (Thermo Fisher Scientific).

Nascent transcripts and chromatin-associated RNAs were separated from the nucleoplasm using the protocol described by ^57^. cDNA was prepared with the use of random-hexamer primers (Thermo Fisher Scientific).

### RT-qPCR

RT-qPCR experiments were performed with Power SYBRR® Green PCR Master Mix (Thermo Fisher Scientific) using a 7900HT Fast Real-Time PCR System (Thermo Fisher Scientific) as previously described ^18^. The primers from Supplementary Table 1 were used. Expression levels were calculated using the relative quantification (2-ΔCt), while the fold change was calculated using the 2-ΔΔCt method ^56^. The mRNA fragments of glyceraldehyde-3-phosphate dehydrogenase (GAPDH, *At1g13440*) were amplified and detected simultaneously as a reference gene.

The *A. thaliana* pri-miRNA expression platform mirEX 2.0 was applied according to ^56^. Each RT-qPCR was performed independently for three biological replicates. All results were analyzed using SDS 2.4 software (Thermo Fisher Scientific) and Microsoft Excel. Error bars were calculated using the SD Function in Microsoft Excel software. The statistical significance of the results presented was estimated using Student’s t test at three significance levels: *P < 0.05, **P < 0.01 and ***P < 0.001.

### Small RNA sequencing

Total RNA was isolated using the Direct-zol™ RNA kit (Zymo Research). RNA was quantified using a Qubit RNA Assay Kit (Life Technologies), and integrity was confirmed on an Agilent Bioanalyzer 2100 system. A total of 10 µg of each RNA sample was separated by electrophoresis on a 15% polyacrylamide 8 M urea gel in 1X TBE buffer. Small RNA fractions were cut out and purified from the gel. Libraries were prepared using the TruSeq Small RNA Library Preparation Kit (Illumina). Single-end (1×50 bp) sequencing was performed at Fasteris, Geneva, Switzerland on a HiSeq 4000 platform. Adapter sequences were removed from raw reads with FASTX-Toolkit (fastx clipper). The clean reads were mapped to all mature *A. thaliana* miRNAs found in miRBase (release 22) using countreads_mirna.pl script ^58,59^. The script was applied to each fastq file for every biological replicate. Statistical analysis was performed with the DESeq2 R package ^60^.

### MicroRNA processing precision calculation

The clean reads from small RNA sequencing were mapped to the *A. thaliana* TAIR10 genome using Rsubread ^61^. The FeatureCount function from the RSubread package was used to obtain the number of reads of precisely processed miRNA (fracOverlapFeature = 1 and fracOverlap = 1 parameters) and imprecisely processed miRNA (fracOverlapFeature = 0 and fracOverlap = 0 parameters). In both cases, only uniquely mapped reads were counted (countMultiMappingReads = FALSE parameter). An annotation file from miRBase (release 22) was used for the mature microRNA coordinates. The processing precision value represents the ratio between precisely and imprecisely processed miRNAs.

### ChIP

Chromatin immunoprecipitation was performed as described ^62^ with IP buffer prepared as described ^63^. Chromatin was sonicated at 4 °C with a Diagenode Bioruptor Pico for ∼15 min (30 s on/30 s off) to obtain 250-500 bp DNA fragments. Antibodies against total RNAPII (Abcam ab817, 5 µg/IP), DCL1 (Agrisera AS19 4307, 10 µg/IP), and SE (Agrisera AS09 532A - 10µg/IP) were used with Dynabeads Protein G (Thermo Fisher Scientific). For decrosslinking and DNA isolation, samples were treated with Proteinase K (Thermo Fisher Scientific) for 6 h at 55 °C followed by purification with a Qiaquick PCR Kit (Qiagen). Libraries were prepared using a MicroPlex Library Preparation Kit (Diagenode) and sequenced on the NextSeq platform.

### Pull-down assay

The *Escherichia coli* strain BL21-CodonPlus(De3)-RIL was transformed with pMal-derived plasmids encoding SE (full length or Δ681-720 aa) or AtPRP40b fused with maltose-binding protein and 6xHis (MBP-6xHis-SE/AtPRP40b). Overexpression was performed as follows: cells were grown for 16 h at 20 °C after induction by 0.4 mM isopropyl β-D-1-thiogalactopyranoside (IPTG) and then harvested and sonicated (6 cycles of 45 s ON and 60 s OFF on ice) in lysis buffer (50 mM Tris–HCl pH 7.5, 300 mM NaCl, 10 mM imidazole, 5 mM β-mercaptoethanol, 0.5% Triton X-100, Roche Complete Mini EDTA-free protease inhibitor tablets (Sigma-Aldrich)). After sonication, lysates were centrifuged for 45 min at 8 000 × g at 4 °C, and the supernatants containing the protein extract were collected. Proteins were purified with HisPur™ Ni-NTA Resin (Thermo Fisher Scientific), and MBP was cleaved off by TEV protease during overnight incubation in dialysis buffer (50 mM Tris–HCl pH=7.5, 300 mM NaCl, 10 mM imidazole, 5 mM β-mercaptoethanol) at 4 °C. The TEV protease and MBP were removed in the additional purification step with the use of HisPur™ Ni-NTA Resin (Thermo Fisher Scientific). Next, SE variants and AtPRP40b were purified by size exclusion chromatography.

The biotinylated CTD peptides were synthetized by Thermo Fisher Scientific. For the pull-down experiment, Streptavidin MagneSphere® Paramagnetic Particles (Promega) were washed three times with Buffer A (PBS pH 8.3, 5% glycerol, 1 mM DTT, 0.03% NP-40, Pierce™ Phosphatase Inhibitor Mini Tablets (Thermo Fisher Scientific)) and incubated with 2 µg of un, Ser5-, or Ser2-phosporylated CTD peptide for 2 h at 4 °C in buffer A. Next, streptavidin particles with immobilized peptides were washed three times with buffer B (PBS pH 8.3, 5% glycerol, 1 mM DTT, 0.1% NP-40, Pierce™ Phosphatase Inhibitor Mini Tablets (Thermo Fisher Scientific), Roche Complete Mini EDTA-free protease inhibitor tablets (Sigma-Aldrich) and incubated with 2 µg of SE variant, AtPRP40b or both for 1.5 h at 4 °C. Streptavidin particles were then washed five times with buffer B, and immobilized proteins were eluted with 3x Laemmli sample buffer (150 mM Tris-HCl pH 6.8, 150 mM DTT, 2% β-mercaptoethanol, 6% SDS, 0.03% bromophenol blue, 30% glycerol) at 25 °C on a thermomixer (350 rpm). Protein samples were separated in a 10% sodium dodecyl sulfate-polyacrylamide gel, transferred to PVDF membranes and detected by Western blots. The following antibodies were used: anti-PRP40b (AS14 2785, Agrisera; 1: 10,000), anti-His (sc-8036, Santa Cruz Biotechnology; 1: 1000), anti-rabbit (AS09 602, Agrisera; 1: 20,000), and anti-mouse (sc-2005, Santa Cruz Biotechnology, 1: 10,000). Input represents 1/10 of the protein sample.

### Western blot

Thirty micrograms of *A. thaliana* whole leaf extract (extraction buffer: 100 mM Tris, 10% glycerol, 5 mM EDTA, 5 mM EGTA, 0.15 M NaCl, 0.75% Triton X-100, 0.05% SDS, and 1 mM DTT) was resolved in a 10% denaturing gel, transferred to PVDF membranes and detected by Western blotting. The following antibodies were used: anti-AtPRP40b (AS14 2785, Agrisera; 1: 10,000), anti-DCL1 (AS19 4307; Agrisera; 1:100), anti-CBP80 (AS09 531; Agrisera: 1: 2000), anti-SE (AS09 532A; Agrisera; 1:2000), anti-HYL1 (AS06 136; Agrisera: 1:1000), anti-actin (691001, MP Biomedicals; 1: 1000), anti-rabbit (AS09 602, Agrisera; 1: 20,000), and anti-mouse (sc-2005, Santa Cruz Biotechnology, 1: 10,000).

### Immunolabeling and FISH

The experiment was performed on isolated nuclei of 35-day-old A. *thaliana* leaves. The leaves were fixed in 4% paraformaldehyde in phosphate-buffered saline (PBS, pH 7.2) for 20 min and washed in 10 mM Tris-HCl (pH 7.5). Nuclei isolation was performed according to the method described in ^64^. The nuclei were permeabilized with PBS+0.1% Triton X-100 for 10 min. The following primary antibodies were used: anti-SE (AS09 532A; Agrisera, 1:100) anti-PRP40b (AS14 2785; Agrisera; 1:100), anti-RNAPII-CTD-Ser5 (3E8; Chromotek; 1:100), anti-RNAPII-CTD-Ser2 (3E10; Chromotek; 1:100) and applied according to ^65^. For the localization of GFP-SE we used mouse antibodies targeting GFP (ab1218; Abcam; 1:100). Primary antibody incubation (in 0.01% acetylated BSA in PBS) was performed in a humidified chamber overnight at 11 °C. After PBS washes, the slides were incubated with the following secondary antibodies: anti-rabbit Alexa Fluor plus 555 (A32732; Thermo Fisher Scientific; 1:200) and anti-rat Alexa Fluor 488 (A-11006; Thermo Fisher Scientific; 1:200) or anti-mouse Alexa Fluor Plus 488, (A32723; Thermo Fisher Scientific; 1:200). The secondary antibodies were diluted in PBS+0.01% acetylated BSA and incubated at 37 °C in a humidified chamber for 1 h.

In double-labeling FISH-immunofluorescence reactions (U1 snRNA + AtPRP40b protein), the in situ hybridization method always preceded the immunocytochemical method. Prior to FISH, the nuclei were permeabilized with PBS+0.1% Triton X-100. The probe targeting U1 snRNA was labeled at the 5′ end with digoxigenin and was resuspended in hybridization buffer (30%, v/v, formamide, 4× SSC, 5× Denhardt’s buffer (0.1% Ficoll 400, 0.1% polyvinylpyrrolidone, 0.1% bovine serum albumin), 1 mM EDTA, and 50 mM phosphate buffer) at a concentration of 50 pmol/ml. Hybridization was performed overnight at 28 °C. Digoxygenin (DIG) probes were detected after hybridization using mouse anti-DIG (11333062910; Merck) and anti-mouse Alexa Fluor 488 (A-11001; Thermo Fisher Scientific, 1:200) antibodies in 0.01% acetylated BSA in PBS.

The slides were stained for DNA detection with Hoechst 33342 (Life Technology) and mounted in ProLong Gold antifade reagent (Life Technologies, P36934).

Correlation analysis was performed with Pearson’s correlation coefficient, Spearman’s rank correlation and the ICQ value. We also used Colocalization Colormap according to ^66^. The statistical analysis was performed using Fiji plugins: coloc2 and Colocalization Colormap ^66,67^. The obtained results were analyzed by Student’s t test, *P <0.001.

### Proximity ligation assay (PLA)

PLA detection was performed using a Duolink In Situ Orange Kit (Merck) according to the manufacturer’s protocol. Prior to the method, the nuclei were treated with PBS buffer containing 0.1% Triton X-100 and then incubated with Duolink blocking solution at 37 °C in a humidified chamber for 60 min. After washing, a 2-stage protocol was applied with the following antibodies: primary rabbit antibodies recognizing SE (AS09 532A, Agrisera, 1: 100) and AtPRP40b (AS14 2785, Agrisera, 1:100), rat antibodies for the detection of phosphorylated RNAPII (serine 5 and serine 2) (Chromotek, 1: 100) and secondary goat anti-rat Alexa Fluor 488 antibodies (A-11006, Thermo Fisher Scientific, 1: 200). The antibodies were diluted in PBS buffer containing 0.05% acetylated BSA, and the incubation was performed overnight at 10 °C (primary antibodies) or at 37 °C for 2 h. After incubation, the nuclei were washed with wash buffer A and subjected to incubation with the Duolink anti-rabbit PLA-plus probe and the Duolink anti-goat PLA-minus probe in Duolink antibody diluent (diluted 1:40) at 37 °C for 1 h. Next, after washing, the slides were incubated with the ligation mix containing ligase at 37 °C for 30 min. Furthermore, amplification buffer containing polymerase was applied. The amplification reaction was performed for 100 min at 37 °C. This type of 2-stage protocol allowed to locate both the total pool of RNAPII (green fluorescence) as well as those that are associated with SE or AtPRP40b (red spots of fluorescence). The slides were stained for DNA detection with Hoechst 33342 (Life Technology) and mounted in ProLong Gold antifade reagent (Life Technologies, P36934).

### Microscopy

The obtained results were registered with a Leica SP8 confocal microscope using a diode 405 laser, an argon/ion laser with a wavelength of 488 nm and a diode laser DPSS 561 that emitted light with a wavelength of 561 nm. For an optimized pinhole, a long exposure time (200 kHz) and 63X (numerical aperture, 1.4) Plan Apochromat DIC H oil immersion lens were used. Images were collected sequentially in blue (Hoechst 33342), green (Alexa 488 fluorescence) and red (Alexa 555, PLA Orange) channels. For bleed-through analysis and control experiments, Leica SP8 software was used.

## Data availability

The data reported in this paper have been deposited in the NCBI GEO database, https://www.ncbi.nlm.nih.gov/geo (accession no. GSE187461).

