## Supplementary figures 1-13 for "Chromatin-associated microprocessor assembly is regulated by PRP40, the U1 snRNP auxiliary protein"

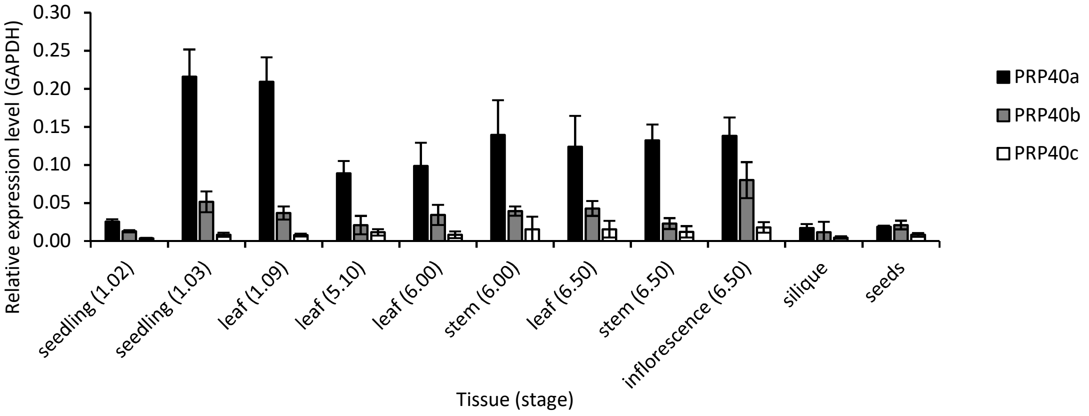


**Extended Data Fig. 1 AtPRP40 proteins are differentially expressed during *A. thaliana* development**
Quantitative RT-PCR showing differences in the expression of the *AtPRP40A*, *B* and *C* genes in different Arabidopsis tissues and growth stages [68].

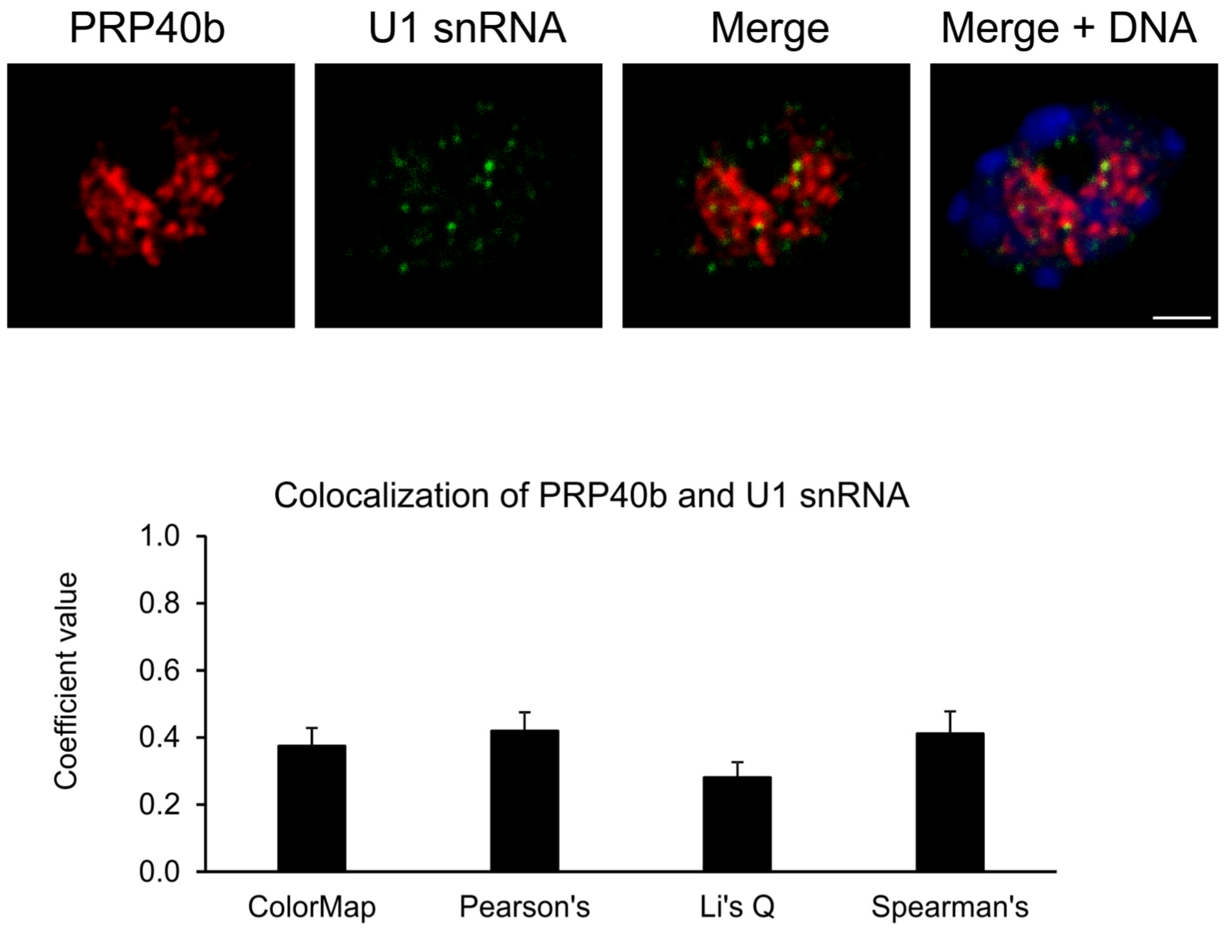


**Extended Data Fig. 2 AtPRP40b is weakly colocalized with U1 snRNA**
Colocalization of AtPRP40b (first image, red signals) and U1 snRNA (second image, green signals) in the cell nucleus. DNA was stained with Hoechst (blue). Scale bar = 5 µm (*upper panel*). Colocalization coefficient values of the colocalization of AtPRP40b and U1 snRNA are shown below the images (*lower panel*).

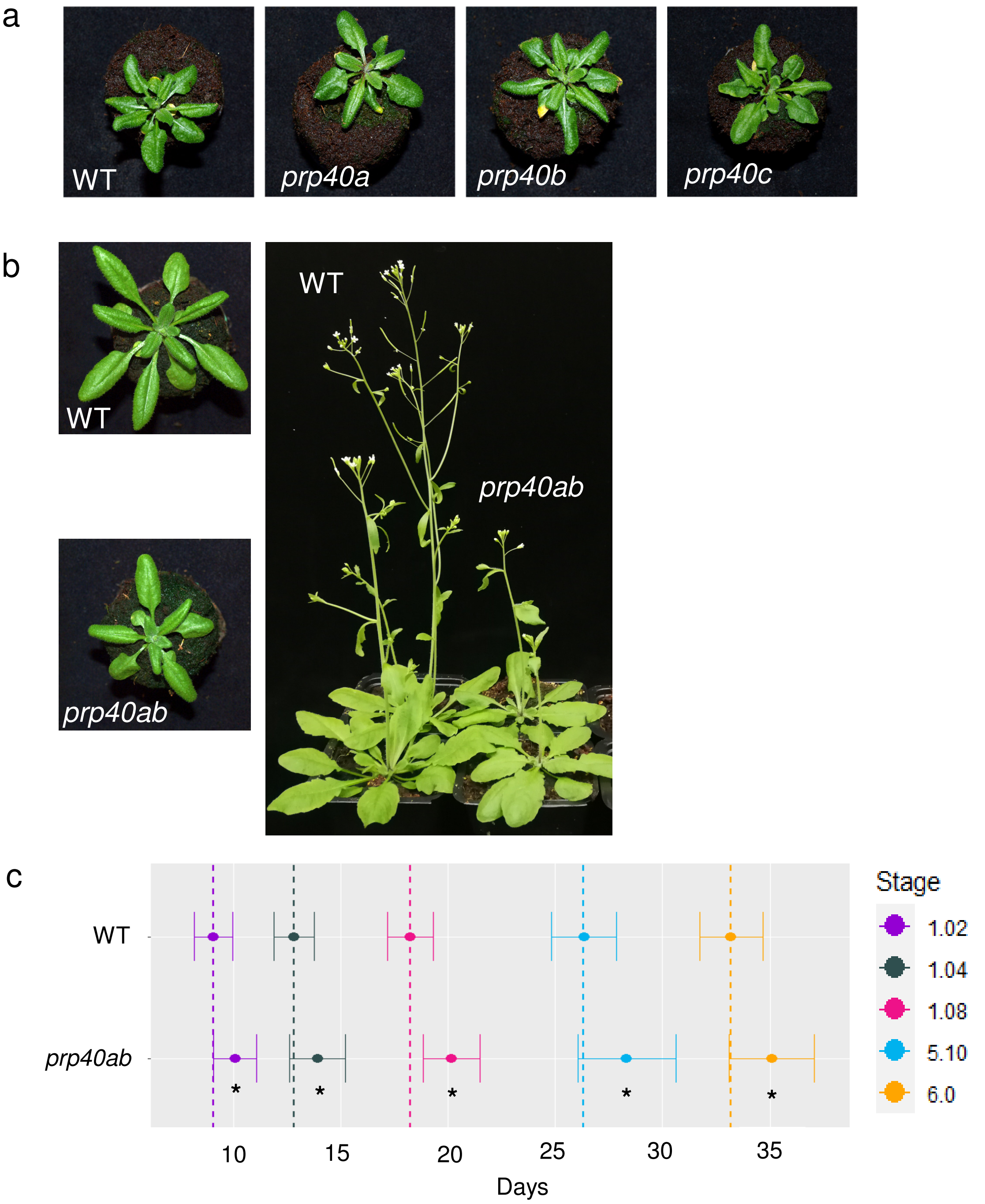


**Extended Data Fig. 3 AtPRP40 is important for *A. thaliana* development**
**a,** Phenotypes of the *prp40a, prp40b* and *prp40c* single mutants. **b,** Phenotype of the *prp40ab* double mutant. **c,** Growth analysis of the *prp40ab* mutant according to [68]

a

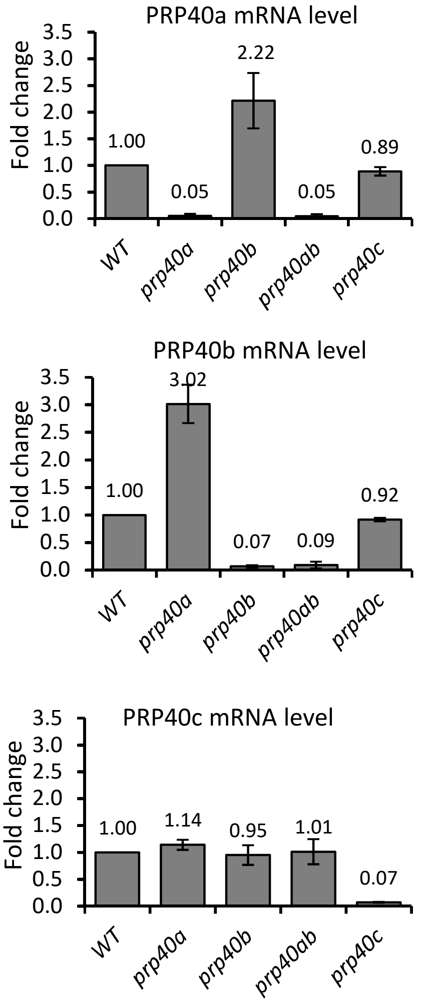


b

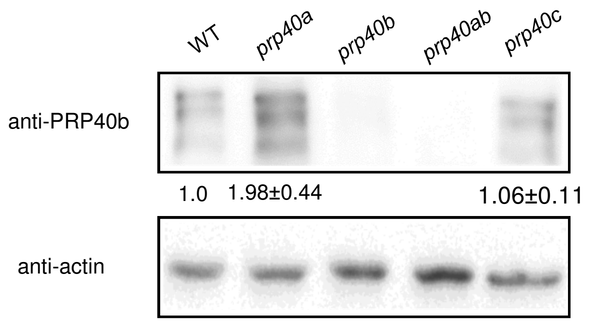


**Extended Data Fig. 4 Redundant role of the AtPRP40a and b proteins**

**a,** Quantitative RT-PCR showing the expression of AtPRP40 genes in the single and double AtPRP40 mutants. **b,** Western blot showing AtPRP40b protein levels in the single and double AtPRP40 mutants.

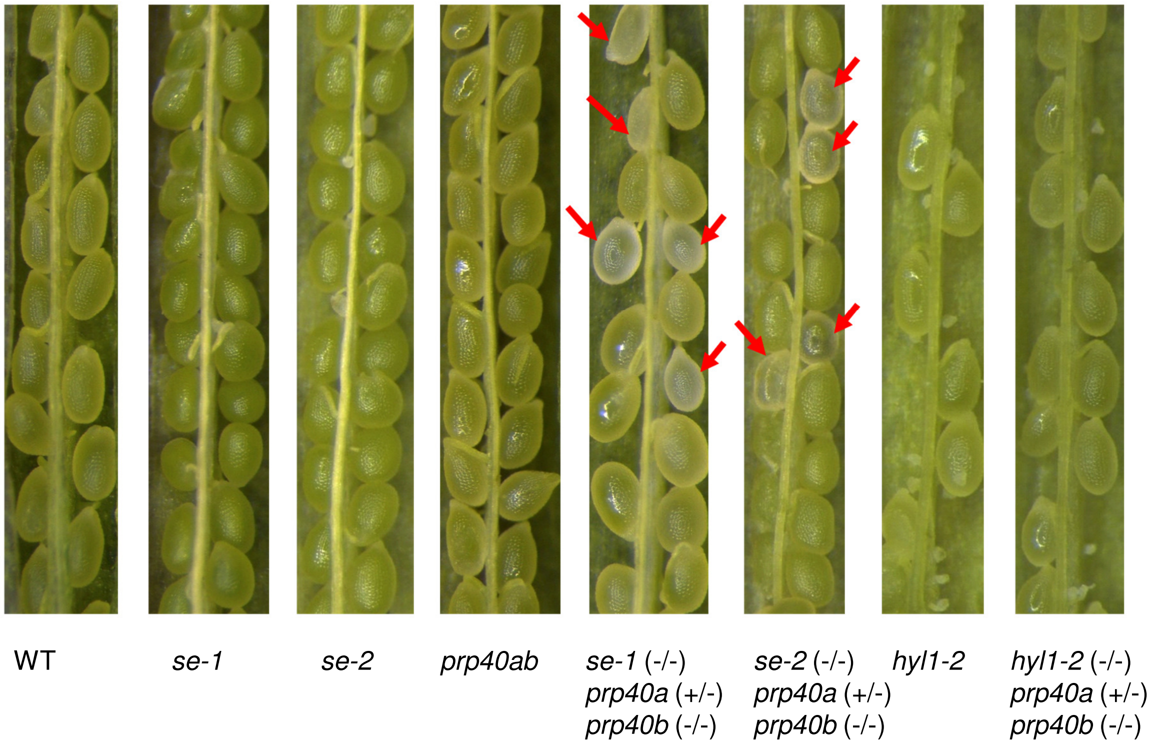


| Seeds | WT | *se-1* | *se-2* | *prp40ab* | *se-1xprp40ab* | *se-2xprp40ab* | *hyl1-2* | *hyl1-2xprp40ab* |
|---|---|---|---|---|---|---|---|---|
| Counted | 188 | 215 | 99 | 205 | 212 | 283 | 35 | 209 |
| % of abnormal | 2.75 | 6.56 | 2.08 | 7.75 | 25.69 | 26.37 | 0.98 | 3.26 |
| SD | 4.24 | 5.57 | 2.95 | 5.37 | 9.31 | 7.23 | 0.05 | 6.30 |

**Extended Data Fig. 5 The crosstalk between SE and AtPRP40 is crucial for plant development** Abnormal, developing seeds are indicated by red arrows. The number and percentage of abnormal seeds for each genotype are presented in the table. SD indicates standard deviation.

**a**

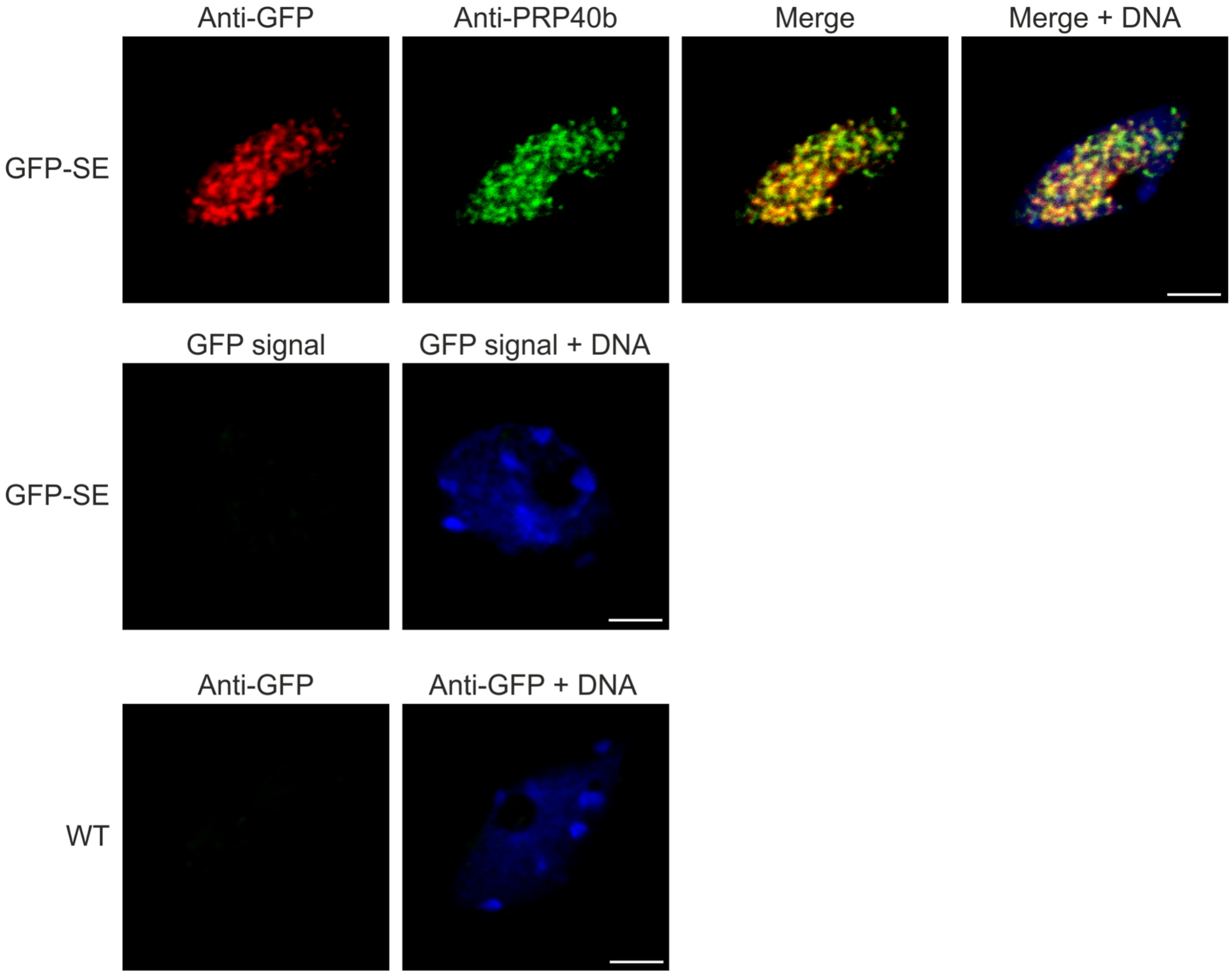


**b**

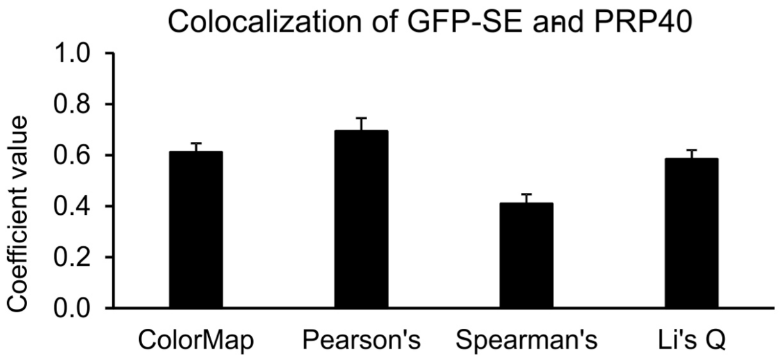


**Extended Data Fig. 6 AtPRP40b colocalizes with SE**

**a**, Colocalization of SE (first image, red signals) and AtPRP40b (second image, green signals) in the cell nucleus of the *A. thaliana* transgenic line expressing GFP-SE. "GFP signal" indicates the signal recorded with the 488 nm laser on a fixed sample prepared without the use of any antibodies. DNA was stained with Hoechst (blue). Scale bar = 5 µm. **b**, Coefficient values for colocalization of GFP-SE and PRP40b in the cell nucleus.

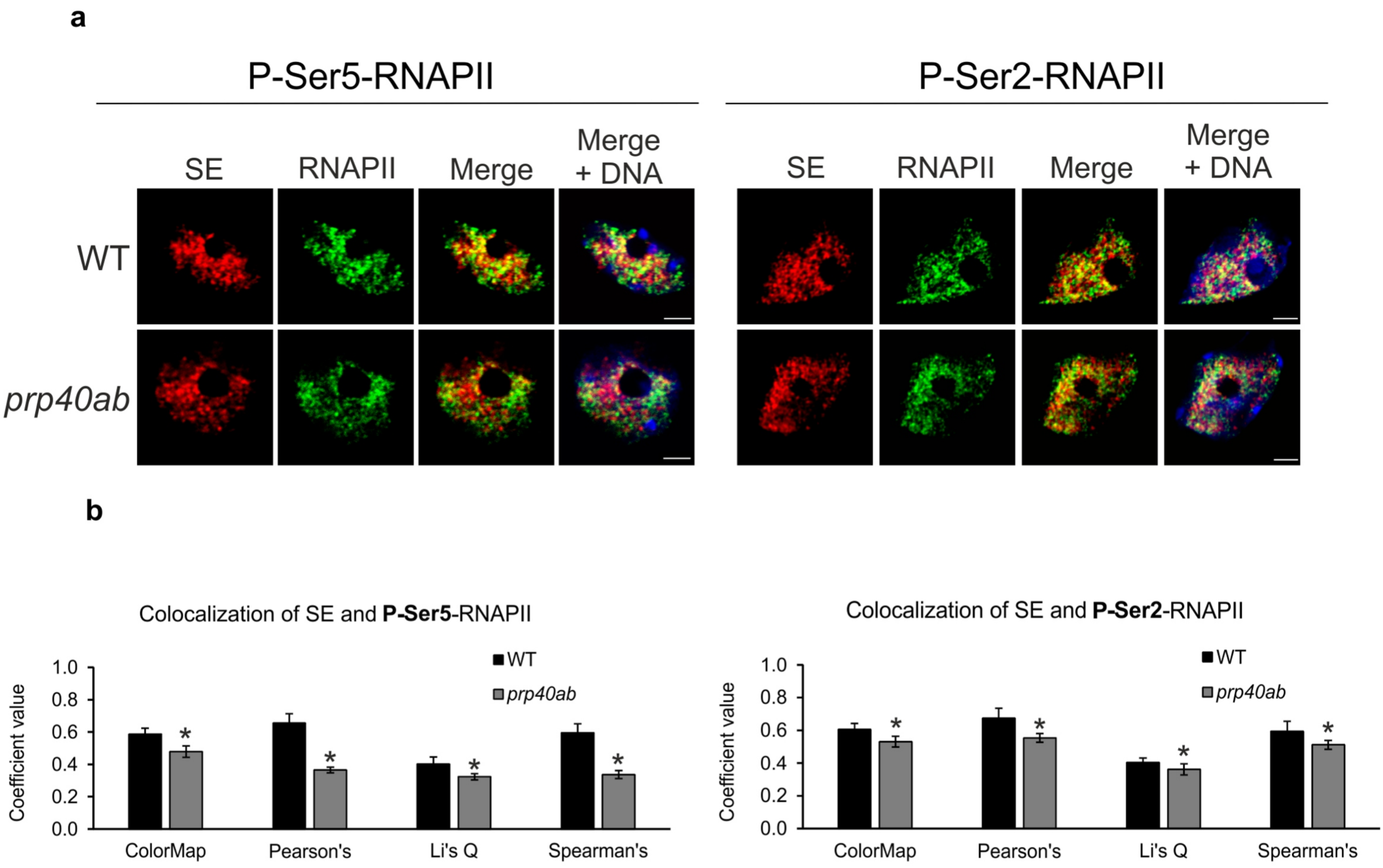


**Extended Data Fig. 7 AtPRP40 regulates the colocalization of SE and RNAPII**
**a**, Colocalization of SE (first column, red signals) and RNAPII phosphorylated at CTD Ser5 (P-Ser5-RNAPII) or Ser2 (P-Ser2-RNAPII) (second column, green signals) in WT and *prp40ab* in the cell nucleus. DNA was stained with Hoechst (blue). Scale bar = 5 μm. **b**, Coefficient values for the colocalization of SE and RNAPII CTD phosphorylated at CTD Ser5 (P-Ser5-) or Ser2 (P-Ser2-) in WT and *prp40ab* cell nuclei. * $P < 0.001$.

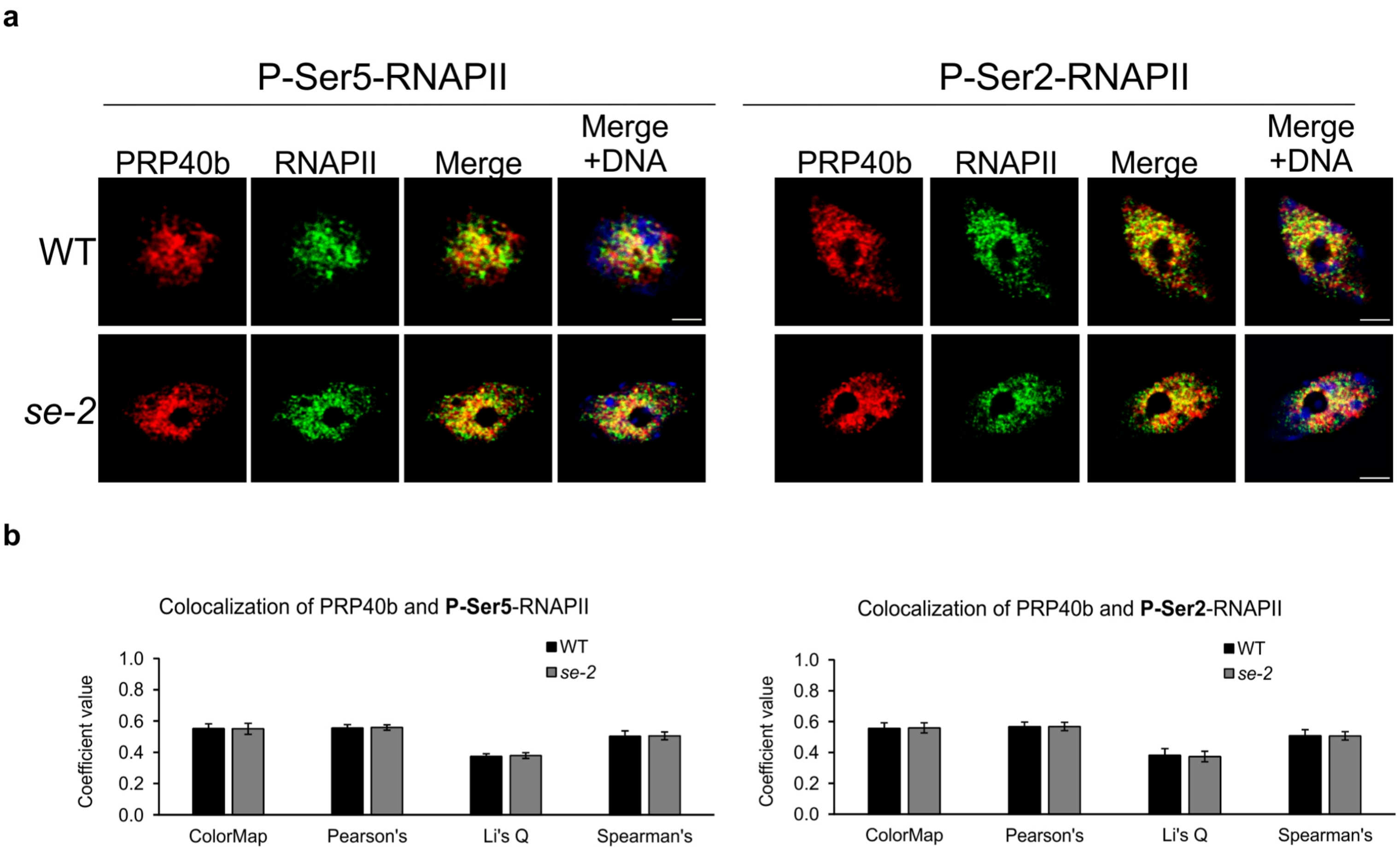


**Extended Data Fig. 8 Colocalization of AtPRP40b and RNAPII in the cell nucleus is not regulated by SE**
**a**, Colocalization of AtPRP40b (first columns, red signals) and RNAPII phosphorylated at CTD Ser5 (P-Ser5-RNAPII) or Ser2 (P-Ser2-RNAPII) (second columns, green signals) in WT and *se-2* cell nuclei. DNA was stained with Hoechst (blue). Scale bar = 5 μm. **b**, Coefficient values for the colocalization of AtPRP40b and RNAPII CTD phosphorylated at CTD Ser5 (P-Ser5-) or Ser2 (P-Ser2-) in WT and *se-2* cell nuclei. * $P < 0.001$.

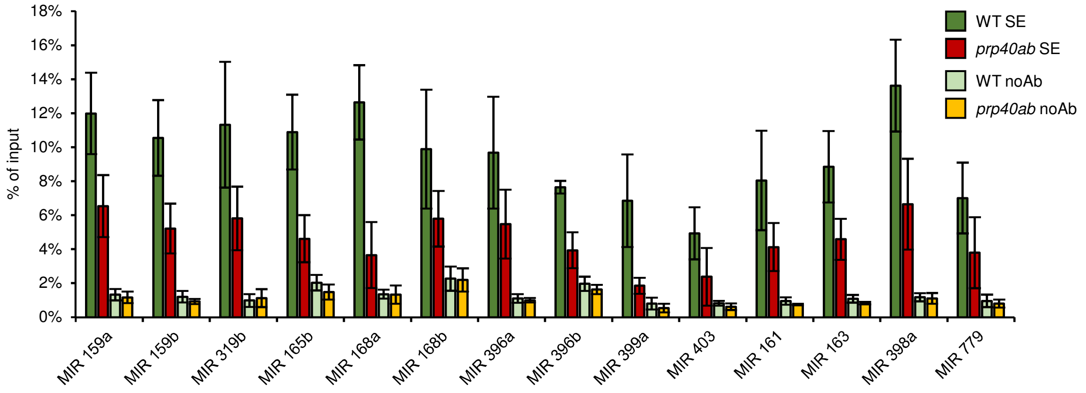


**Extended Data Fig. 9 AtPRP40 is required for the proper accumulation of SE on miRNA genes**
Quantitative ChIP-PCR showing the level of SE on miRNA genes in the *prp40ab* mutant (n=3).

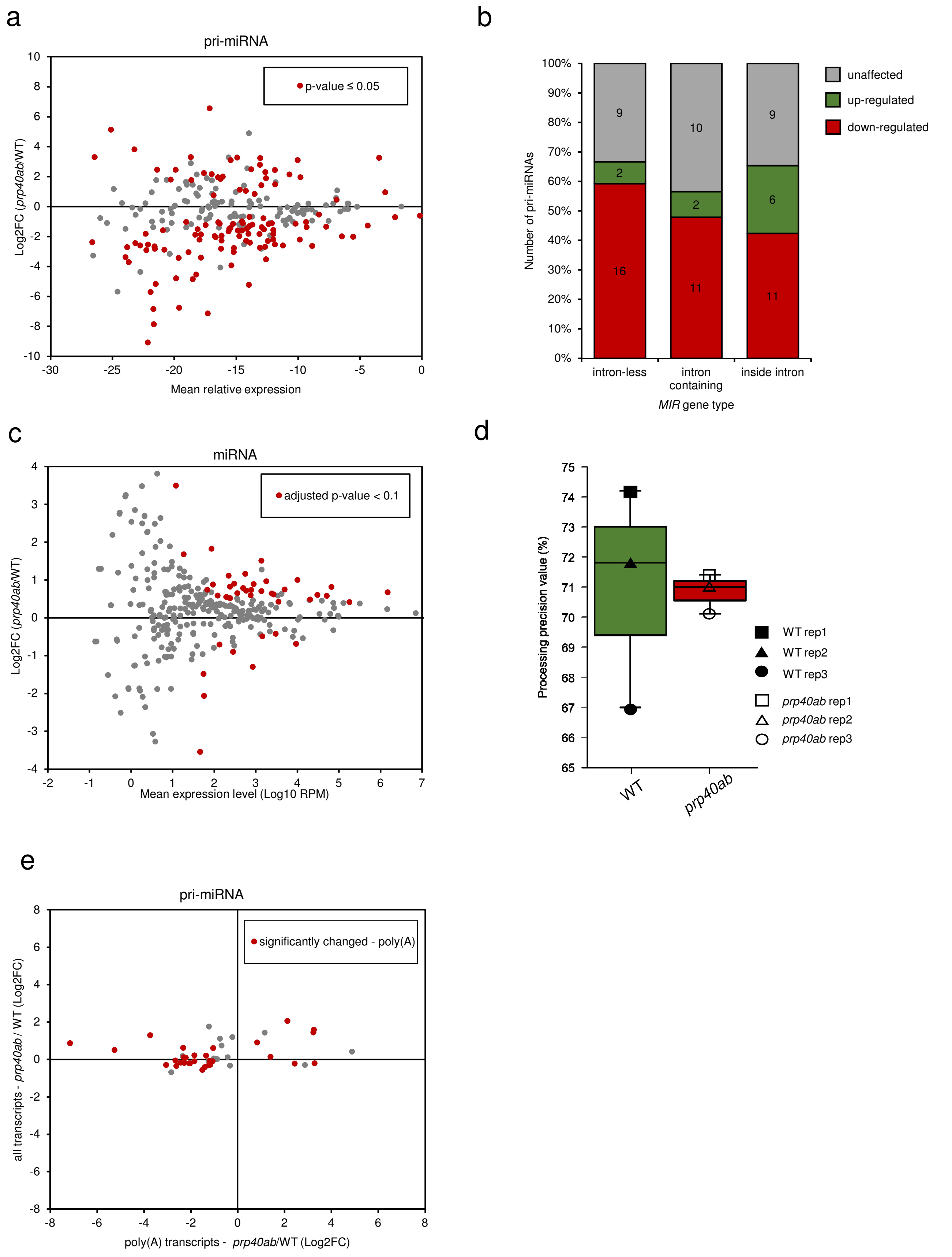


**Extended Data Fig. 10 The effect of AtPRP40 on miRNA biogenesis**

**a,** MA plot showing a relation between the fold change (*prp40ab*/WT) of polyadenylated pri-miRNA and its mean relative expression. **b,** Number of changed polyadenylated pri-miRNAs in *prp40ab* depending on the miRNA gene type. **c,** MA plot showing a relation between the fold change (*prp40ab*/WT) of miRNA and its mean relative expression. **d,** Box plot showing **the** precision of miRNA processing in *prp40ab* plants compared wild-type plants. **e,** Fold change (*prp40ab*/WT) comparison for polyadenylated and total *MIR* transcripts.

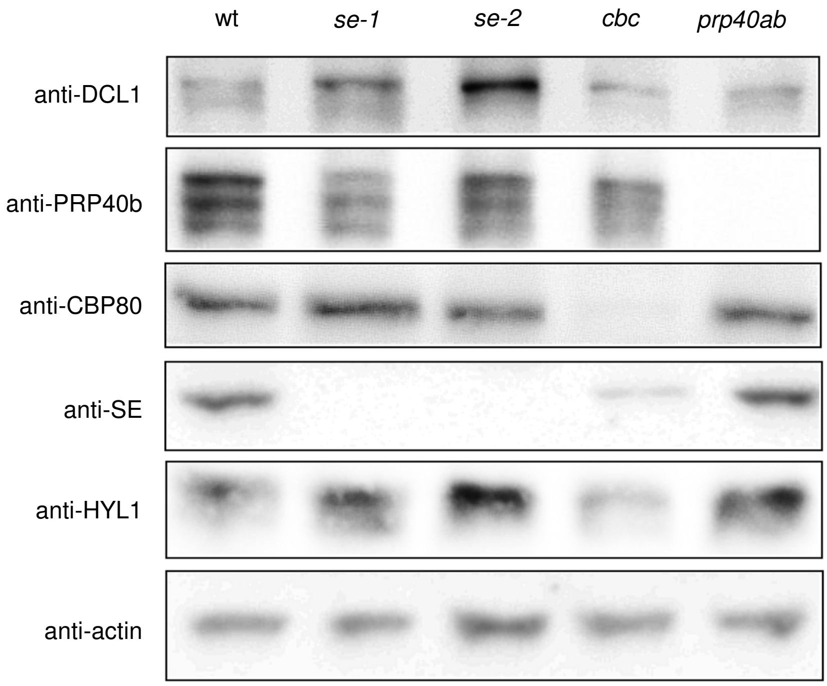


**Extended Data Fig. 11 The levels of miRNA biogenesis-related proteins in *prp40ab***
DCL1, SE, HYL1, CBP80 and PRP40b levels in the *se-1*, *se-2*, *cbp20xcbp80* and *prp40ab* mutants determined by Western blots. The actin level was used as a loading control. The experiments were performed with three replicates.

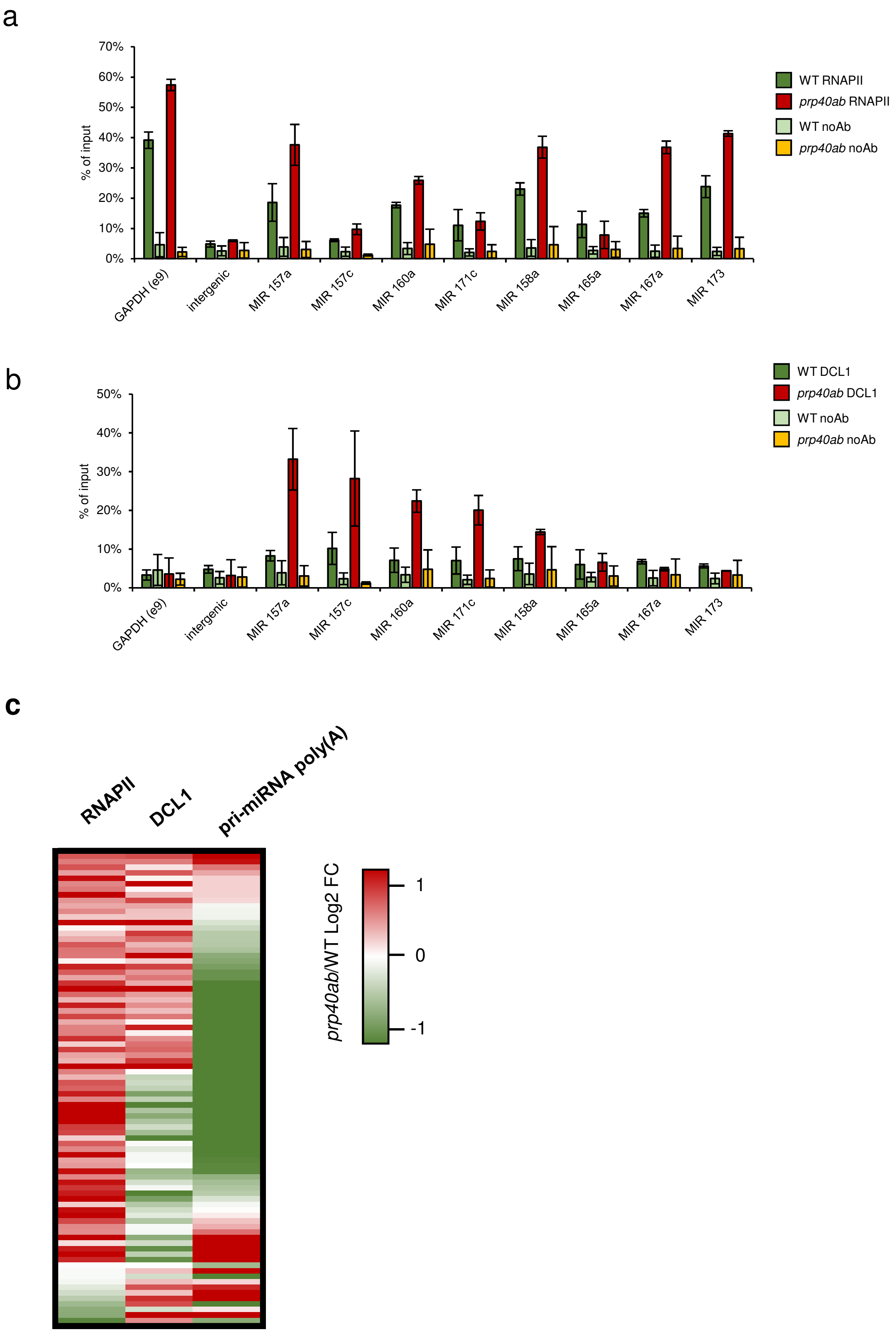


**Extended Data Fig. 12 RNAPII and DCL1 distributions on miRNA genes are both affected in the *prp40ab* mutant**

**a,** Quantitative ChIP-PCR showing changes in the RNAPII distribution on miRNA genes in the *prp40ab* mutant (n=2). **b,** Quantitative ChIP-PCR showing changes in the DCL1 distribution on miRNA genes in the *prp40ab* mutant (n=2). **c,** Heatmap based on RNAPII ChIPseq, DCL1 ChIPseq and quantitative RT-PCR data for miRNA precursors. Only precursors with data available from three experiments are shown.

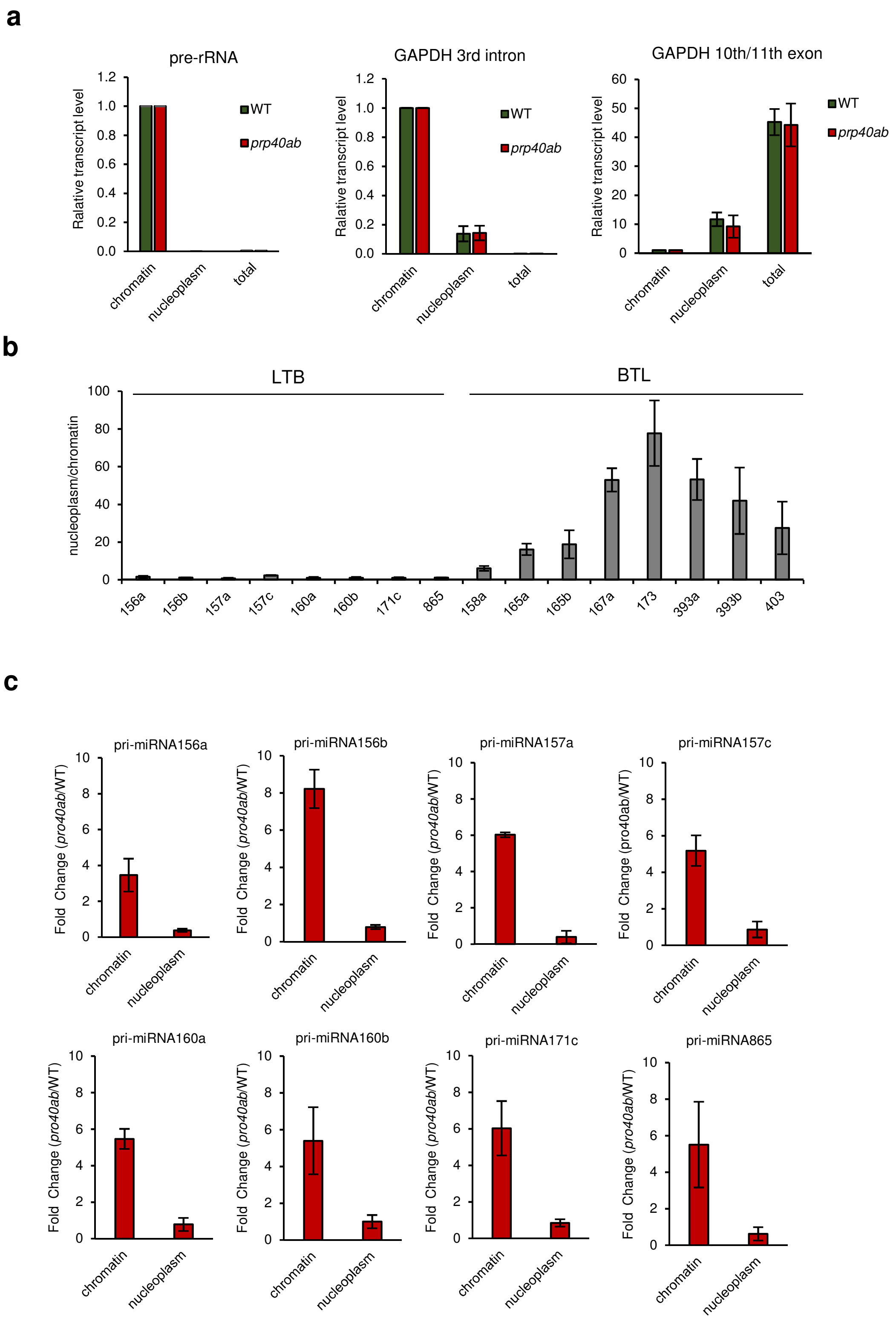


**Extended Data Fig. 13 The distribution of miRNA gene transcripts between chromatin and the nucleoplasm is affected in the *prp40ab* mutant**

**a,** Quantitative RT-PCR showing changes in the distribution of control transcripts depending on the tested fractions (n=3). **b,** Quantitative RT-PCR showing changes in the nucleoplasm/chromatin ratio depending on pri-miRNA processing type (LTB – loop-to-base, BTL – base-to-loop) (n=3) in WT plants. **c,** Loop-to-base-type pri-miRNA fold change (*prp40ab*/WT) for chromatin and nucleoplasmic fractions (n=3). **d,** Base-to-loop-type pri-miRNA fold change (*prp40ab*/WT) for chromatin and nucleoplasmic fractions (n=3). **e,** Polyadenylated pri-miRNA fold change (*prp40ab*/WT) for chromatin and nucleoplasmic fractions (n=3).

**d**

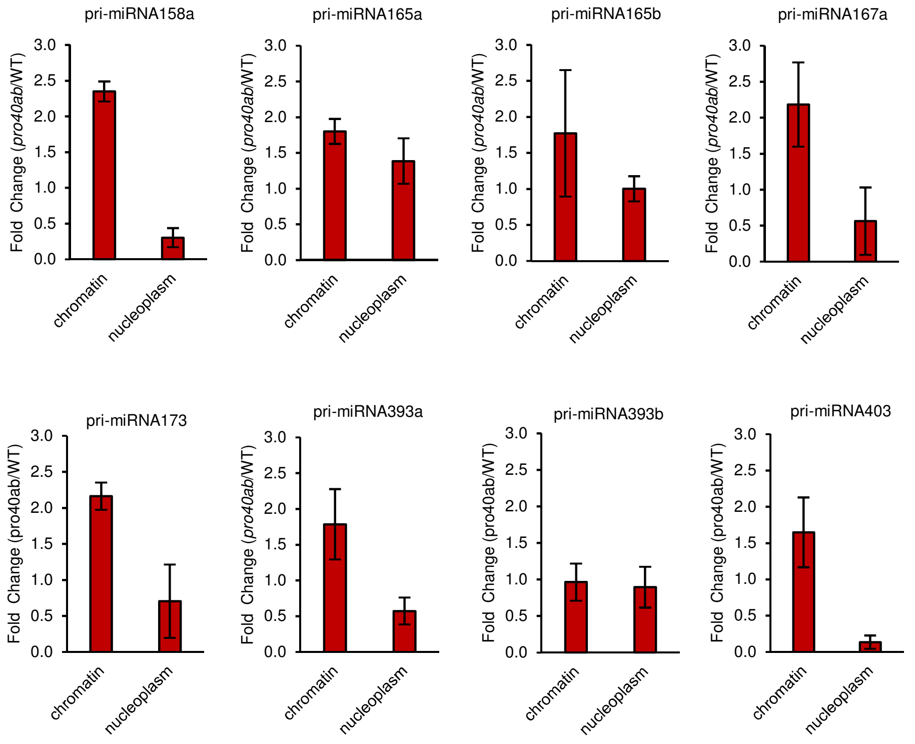


**e**

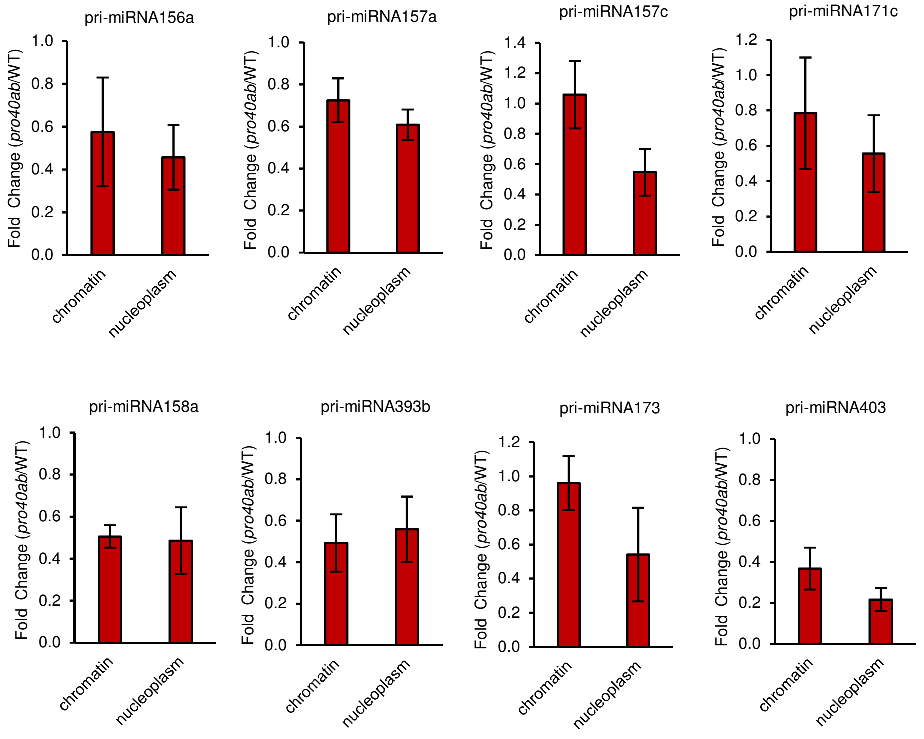


**Extended data Fig 13 Continued.**
