## Supplementary table 1 for "Chromatin-associated microprocessor assembly is regulated by PRP40, the U1 snRNP auxiliary protein"

**Supplementary table 1. Primers used in this study**

| Name | Sequence (5' → 3') |
|---|---|
| F_GAPDH_i3 | TTCGTTTCTTCTTCTCTTCGAT |
| R_GAPDH_i3 | TGATAAACAGCACACACCATCA |
| F_GAPDH_e9 | TGGAAAATTGACCGGAATGT |
| R_GAPDH_e9 | TCGTCGTATGTTGCAGCTTT |
| F_GAPDH_e10/11 | TTGGTGACAACAGGTCAAGCA |
| R_GAPDH_e10/11 | AAACTTGTCGCTCAATGCAATC |
| F_pre_rRNA | GCGAACCAAAGATCACCACT |
| R_pre_rRNA | TTCGTTTGCATGTTCCTTGA |
| F_intergenic | AGTTCAATGGAGAGATGTCGAAATATG |
| R_intergenic | AAGAGGAAAAGAAAGAGATGGAGAGA |
| F_PRP40a | ACACTTGGCCTGTTCCTGTT |
| R_PRP40a | TGCGGAGTCAAGTTTCCTGG |
| F_PRP40b | AAGCACCGTGATGAGTTCCA |
| R_PRP40b | CCAGAGGAGTTTGACGCAATTG |
| F_PRP40c | CAAGGAATGTGGTTGCAGCC |
| R_PRP40c | CCAAGCGGATGAGAACCAGA |

For pri-/pre-miRNA amplification mirEX platform primers were used [37]
